# Yeast and Mammalian Epsins Use Different Determinants for Localization and Function: Role of Clathrin/AP2/Ubiquitin Binding Motifs and Poly-Glutamine Stretches

**DOI:** 10.1101/2022.08.04.502746

**Authors:** Kayalvizhi Madhivanan, Sneha Subramanian, Lingyan Cao, Debarati Mukherjee, Arpita Sen, Wen-Chieh Hsieh, Claudia B. Hanna, Beibei Wang, Hong Chen, Chris J. Staiger, R. Claudio Aguilar

**Affiliations:** Department of Biological Sciences, Purdue University; Purdue Center for Cancer Research, Purdue University; Vascular Biology Program, Boston Children’s Hospital, Boston, MA; Department of Botany and Plant Pathology, Purdue University, West Lafayette, IN 47907 USA

**Keywords:** Endocytosis, Epsin, Clathrin, Ubiquitin, AP2, Poly-Glutamine

## Abstract

Epsins are endocytic adaptor proteins involved in the internalization of important membrane proteins such as EGFR and Notch ligands. Therefore, this protein family impacts critical signaling pathways and processes such as cell migration and cytokinesis and is ultimately required for embryo development in mammals and cell viability in yeast. Intriguingly, although Epsins are conserved and display similar binding determinants, the process of endocytosis in yeast and mammals exhibit some dramatic mechanistic differences. Therefore, we wondered if the function of Epsins in these organisms are similar and are similarly regulated or they also differ. Since proper and timely localization is needed for function, we determined what elements target Epsins to endocytic sites in yeast vs mammals. Specifically, using a systematic/combinatorial mutagenesis approach we produced a collection of yeast and human Epsin mutated variants that was tested for localization at endocytic sites and for function.

Our results showed that the intrinsically disordered carboxy-terminus holds the major determinants (involved in binding of ubiquitin, AP2, clathrin and EH domain-containing proteins) for proper intracellular localization of different Epsin paralogs and homologs in yeast and mammals, while also having a major impact on function. Importantly, we established hierarchies of carboxy-terminal binding determinants for sustaining Epsin localization which turned to be different for human vs. yeast cells; favoring clathrin and AP2 binding in the former and recognition of cargo and EH domain-containing proteins for the latter. Further, we found evidence in both systems that yeast Epsins also use for localization regions of the protein that were until now of unknown functional relevance, *i.e.,* glutamine-rich sequences. Interestingly, some molecular determinants within the Epsin molecule seem to have functional importance beyond its contribution to localization to endocytic sites. Based on these findings, we propose working models for Epsin function and recruitment to membranes/endocytic sites at different maturation stages.

## INTRODUCTION

Endocytosis is a process that not only is required for the acquisition of nutrients but also for membrane composition control. Importantly, endocytosis contributes to the spatial and temporal regulation of membrane proteins such as plasma membrane receptors. Therefore, endocytic pathways are critical for the regulation of intracellular signaling, ultimately impacting other key cellular processes such as cell proliferation, migration, and cytokinesis. Consequently, understanding the mechanism of action of the intracellular machinery involved in endocytosis is of great importance for our understanding of the patho-physiology of multiple diseases, but also for the design of effective therapeutic approaches (*e.g.,* for the intracellular delivery of therapeutics).

Among the intracellular machinery required for endocytosis, coat-associated adaptors are central components as they are responsible for cargo selection. The Epsin (Epn) family of endocytic adaptors bind Phosphatidylinositol (4, 5) bisphosphate (PI(4,5)P_2_)-enriched membrane regions, the clathrin coat and ubiquitinated cargoes such as EGFR (Epidermal Growth Factor Receptor) and VEGFR2 (Vascular Endothelial Growth Factor 2) (Bertelsen et al., 2011, Kazazic et al., 2009, Sigismund et al., 2005, Pasula et al., 2012). Further, and besides this adaptor function, members of this protein family can also act as endocytic accessory proteins by binding to elements of the endocytic machinery to stabilize the endocytic interaction network. In addition, it has been demonstrated that Epn contributes to the initiation of membrane curvature at sites of nascent endocytosis (Ford et al., 2002) and is one of the endocytic checkpoint proteins that decide if nascent coated pits should continue maturation to the point of vesicle scission or if they should be disassembled (Mettlen et al., 2009). However, and although other proteins within the endocytic network could potentially substitute for loss of Epn function, these proteins are essential for embryo development and cell viability in eukaryotes (Fischer et al., 1997; Tian et al., 2004; Chen et al., 2009; Wendland et al., 1999, Aguilar et al., 2006), likely due to the endocytosis-dependent, Epn-mediated signaling modulation of the critical EGFR, VEGFR2 and Notch signaling pathways (Wang et al., 2006a; Tian et al., 2004; Musse et al., 2012). Epns also play an important role in signaling regulation related to RhoGTPases (Aguilar et al., 2006; Coon et al., 2010), Protease-Activated Receptors (Chen et al., 2011) but also in the processes of cell migration and invasion (Coon et al., 2010, 2011). Importantly, the latter two processes are quite relevant to the observation that gastric and other cancers show upregulation of some Epn paralogs suggesting relevance of this protein family in the disease’s invasive characteristics (Coon et al., 2010, 2011).

In unicellular organisms like *S. cerevisiae* many of the processes in which Epns are involved are necessarily different. Further, although yeast homologs show motifs shared with the higher eukaryote’s Epns, the spatial organization of such determinants within the protein is different and except for the ENTH domain, the overall sequence conservation is low (Sen et al., 2012). In general, substantial differences with metazoans also exist in terms of mechanisms of endocytosis as in yeast clathrin and the emblematic tetrameric adaptor proteins (APs) are not essential (Boettner et al., 2011), the actin cytoskeleton has a more crucial role, with an still uncertain involvement of dynamin-like proteins and marked differences in the overall structural organization of the endocytic pit (Idrissi et al., 2008; Idrissi & Geli 2014).

The goal of the present study was to determine the role of different molecular determinants supporting Epn function in yeast and mammalian cells. Specifically, and since proper and timely localization is important for function, we studied the elements mediating targeting of Epns to endocytic sites and how this affect their role in cargo/vesicle internalization dynamics and specific cellular processes such as cell migration.

Here we show that the mostly intrinsically disordered carboxy-terminus is the major determinant for proper intracellular localization of different Epn paralogs and homologs in yeast and mammals, while also having a major impact on function. Specifically, we established a hierarchy for the requirement of the endocytic determinants found in the Epn’s carboxy-terminus for localization which was different for human vs. yeast cells; favoring clathrin and AP2 binding in the former and suggesting a more important role for the recognition of cargo and EH domain-containing proteins for the latter. Further, both systems displayed evidence of also using regions of the protein for localization that were until now of uncertain functional relevance, *e.g.,* a paralog-specific region in human Epn3 and a subset of glutamine-rich sequences in the yeast homologs. Finally, as hypothesized, overall proper localization was needed for Epn to fulfill the specific functions monitored in this study (endocytic pit maturation and cell migration in mammalian cells; cargo internalization and cytokinesis control in yeast). However, some molecular determinants within the Epn molecule seem to have functional importance beyond its contribution to localization to endocytic sites. These observations suggest that specific Epn functions relied on certain interactions (*e.g.,* mediated by ubiquitin and EH domain-containing protein motifs) in higher eukaryotes and (by glutamine repeats) in yeast.

In addition, and based on these findings, we propose working models for Epn function and recruitment to membranes/endocytic sites at different maturation stages.

This study used a combinatorial, systematic mutagenesis approach to analyze the relative roles of molecular determinants for localization/function. We were able to show not only the relevance of different Epn’s interactions for such readouts, but also highlight differences between the yeast model and human cells. We envision that similar approaches will be also proven to be powerful to understand molecular mechanism of action of many other proteins than like Epn display multi-specific, multi-valent intrinsically disordered regions.

## RESULTS

### The carboxy-terminus contains the major determinants for intracellular localization of the Epn endocytic adaptor

Epns are a conserved family of endocytic adaptors characterized by an ENTH domain and an intrinsically disordered carboxy-terminus (Fig.1A and Sen et al., 2012). While the ENTH domain of Epn is known to bind PI(4,5)P_2_-enriched regions at the plasma membrane, the carboxy-terminus binds to different components of the endocytic machinery including ubiquitinated cargoes, clathrin, AP2 and EH domain-containing proteins such as Eps15 and Intersectin (Fig.1A and Sen et al., 2012). Further, sequence alignment of the three mammalian Epns shows that their ENTH domains are highly conserved, and although the carboxy-terminal sequences are the most divergent, they still show a common organization of endocytic binding motifs (Fig. 1A). However, when compared to the yeast Epns Ent1 and Ent2, marked differences in motif organization and sequence can be observed (Fig.1A and Sen et al., 2012).

**Fig. 1.**
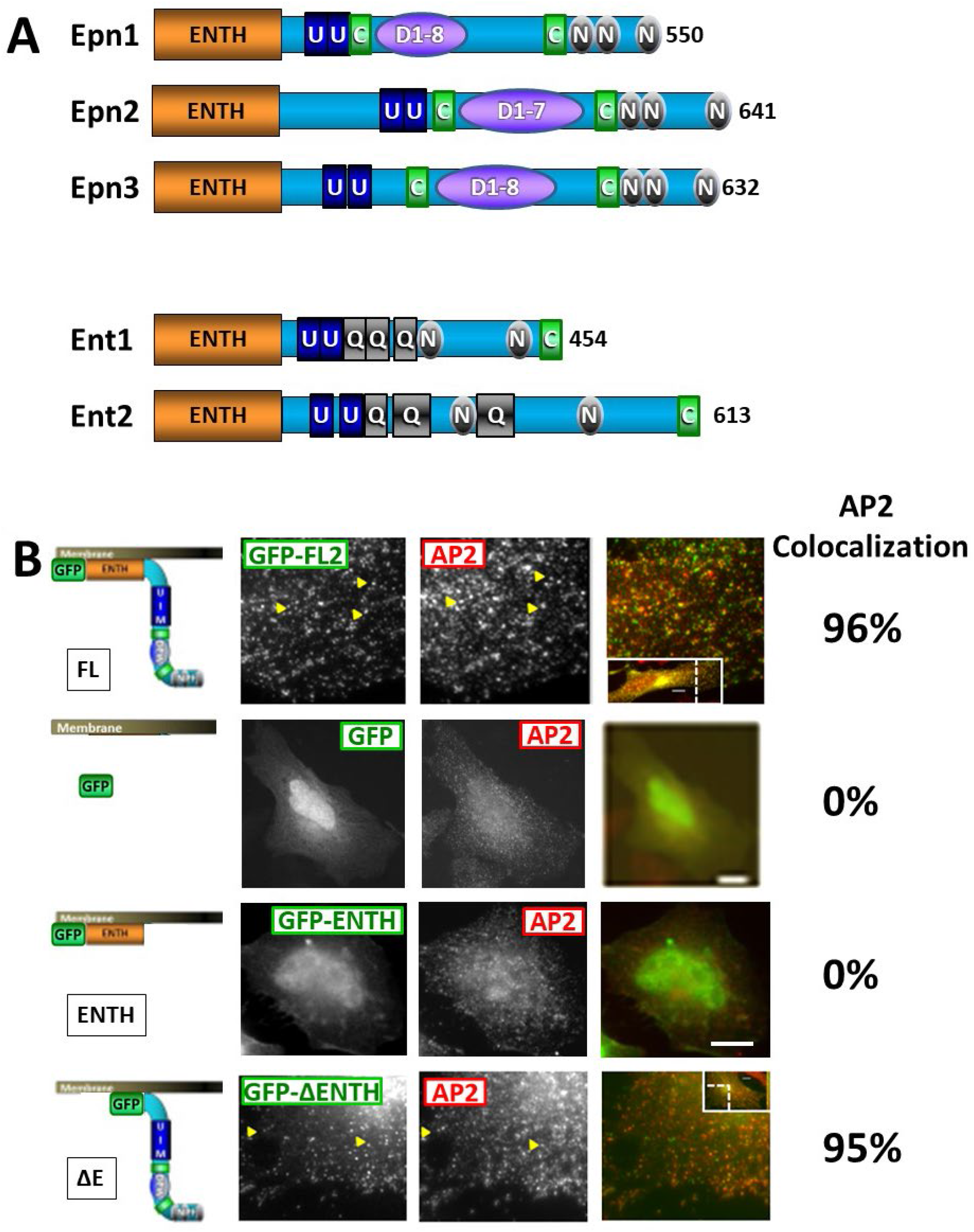
A. Domain/Motif Architecture of human and yeast Epns. Ubiquitin interacting Motifs (U) in blue, Clathrin Binding Motifs (C) in green, AP2 Binding Motifs (D: DPW/DPF and similar EPW/NPW/QPW) in purple and NPFs (N: Asparagine, Proline, Phenylalanine) in grey. Glutamine-rich regions (Q) in yeast Epns Ent1/2 are shown in grey. **B. Colocalization of AP2 with GFP and different truncations of GFP-Epn2** (GFP-FL, GFP-ENTH, GFP-ΔΕ). Results from quantitative analysis is also shown. Arrowheads point to some examples of colocalizing puncta. Scale Bar = 10uM. Please refer to text for details.

In mammals, Epns function as adaptors, accessory and endocytic checkpoint proteins, consequently they are found localized at the plasma membrane in endocytic sites marked by the presence of clathrin and the clathrin-associated, tetrameric adaptor AP2. Since presence at the correct site and proper time is critical for function, we started by testing the putative role of different Epn’s regions and motifs in protein intracellular localization. In fact, although the ENTH domain plays a role in membrane attachment, it is not completely clear if it is sufficient for localization or if there are other major localization determinants of Epn and whether such putative determinants are the same across paralogs and species. To address these questions, we used a systematic mutagenesis approach focusing on mammalian Epn 1, 2 and 3.

We selected Epn2 to start the series given the availability of *EPN2* K.O. cells (see below), although analogous mutants were prepared and tested for Epn1 and 3. Specifically, we first split the Epn2 molecule in two: the N-terminal ENTH domain (ENTH2-GFP) and the complementary carboxy-terminal truncation lacking such domain (ΔENTH2: ΔE2-GFP). We proceeded to express such truncations in HeLa cells and compared their localization to Full-Length Epn2 (FL2-GFP) as well as to isolated GFP (*i.e.,* empty pEGFP vector). Samples were fixed and co-stained by indirect immunofluorescence using specific antibodies against AP2 (see *Materials and Methods*) to test their ability to localize to AP2-positive endocytic sites. Although both AP2 and clathrin are the major markers of clathrin-mediated endocytic (CME) sites, we immunostained only AP2 because it is found *exclusively* at the plasma membrane, while clathrin is also known to be present at endosomes and the trans-golgi network (TGN) (Keyel et al., 2004). In addition to proper localization to endocytic sites, we also evaluated potential mislocalization of these GFP-fusions to the TGN, nucleus and cytoplasm (details on colocalization calculations can be found in *Materials and Methods*).

In agreement with the work of others (Rosenthal et al., 1999), our results showed that FL2-GFP was present in almost every endocytic site marked by AP2, showing a virtual one-to-one (96%) colocalization (Fig.1B, top row). In contrast, free GFP and ENTH2-GFP did not colocalize with AP2. Although ENTH2-GFP showed evidence of some level of membrane association, it lacked enrichment at AP2-sites and was mostly confined to the cytosol and nucleus. Nuclear accumulation is a well-known artifactual feature of free/isolated GFP (Fig.1B, 2^nd^ row) and of GFP-protein fusions where the fused protein lacks strong localization determinants. Therefore, we concluded that the isolated ENTH2 domain (ENTH2-GFP) despite to be expected to bind PI(4,5)P2, provided weak and non-CME specific membrane localization properties (Fig.1B, 3^rd^ row).

Importantly, ΔΕ2-GFP showed a 95% colocalization of with AP2-positive sites. However, ΔE2 also showed partial mislocalization to the cytoplasm. As a whole, these results indicated that the carboxy-terminus of Epn2 contain the major determinants of localization; nevertheless, given that the absence of the ENTH domain led to partial mislocalization to the cytosol, we speculate that it contributes to enhance the specificity and binding to the plasma membrane. The role of the Epn carboxy-terminus in protein localization was conserved across paralogs and also yeast homologs (Supplemental Fig.1).

### Carboxy-terminal determinants involved in the recognition of endocytic proteins contribute to Epn2 localization

Next, we tested the contribution of carboxy-terminal individual motifs to the localization of Epn2. Indeed, as mentioned before, this Epn region contains short determinants for binding to four different components present in endocytic sites: ubiquitinated cargoes (*via* two ubiquitin-interacting motifs, U), clathrin (*via* two clathrin binding motifs, C), AP2 and EH domain-containing proteins (*via* seven DPW/DPW-alike and three NPF tripeptides—D and N, respectively) (Figs.1 and 2).

We proceeded to perform systematic combinatorial mutagenesis of these motifs (Fig. 2) in the context of the FL and ΔΕ constructs of Epn2. As a first step, we mutated all motifs for binding one given component present in endocytic sites according to figure 2 (*e.g.,* both C motifs) and then proceeded to generate all possible combinations of mutated determinants (see also *Materials and Methods*). We named the resulting mutant according to the *motifs left intact* followed by the *paralog identification number*; for example, while UCDN2 represents a construct with WT carboxy-terminal sequence of Epn2 (*i.e.,* all four determinants for binding to ubiquitin, clathrin, AP2 and EH domain containing proteins are unaltered), CDN2 refers to an Epn2 mutant lacking functional U (ubiquitin binding) sequences. Similarly, UCN2 represents Epn2 with its seven D-motifs mutated and CN2 encodes for the protein with its two U and seven D determinants inactivated.

**Fig. 2.**
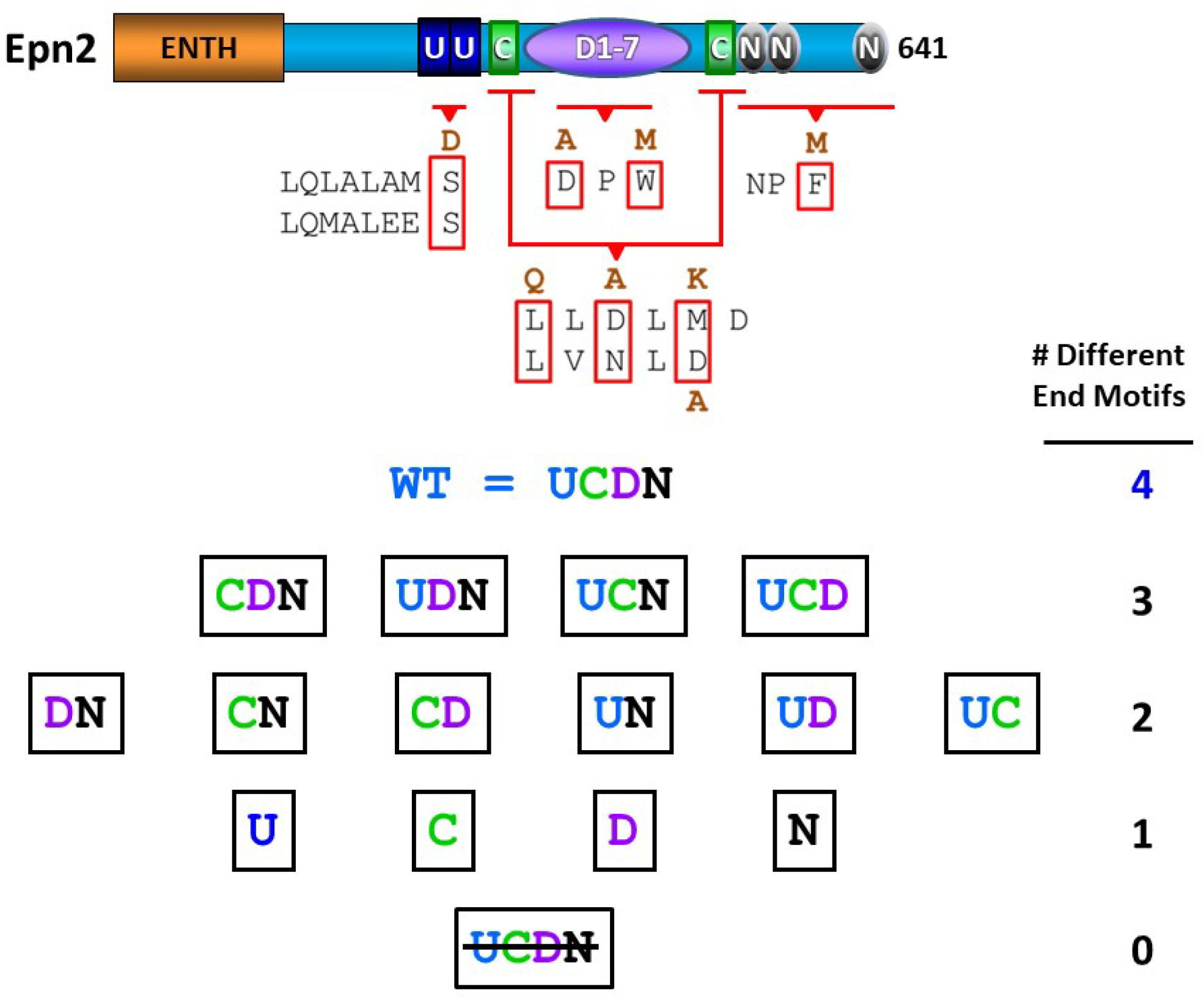
Combinatorial mutational approach used on Epn2. The WT motif sequence along with the mutations introduced are shown. UCDN refers to the WT sequence. Resulting combinatorial mutants are as follow: first row (CDN, UDN, UCN and UCD) refers to mutants with one type of endocytic determinant mutated. The second row (DN, CN, CD, UN, UD, UC) represent mutants with two types of endocytic determinants mutated. Third row (U, C, D, N) refers to mutants with three types of endocytic determinants mutated. represents a mutated variant in which all four types of binding motifs have been altered. Please refer to text for details.

Following this approach and nomenclature, we prepared all four mutated variants displaying three endocytic determinants (*i.e.,* missing only one kind of binding motif: CDN2, UDN2, UCN2 and UCD2), six mutants with two kinds of binding sites (DN2, CN2, CD2, UN2, UD2 and UC2), the four mutants bearing only one type of determinant (U2, C2, D2 and N2) and the one construct having *none* of the 4 binding motifs intact (represented as 2). It should be noted that in the FL-2 variant with all short binding determinants mutated, >90% of the protein sequence still remains unaltered. For full list of constructs used in this study and the nature of the mutations introduced, see Fig.2 and *Materials and Methods*. The stability of mutated variants was assessed by western blotting showing no substantial difference in protein stability (data not shown).

GFP-fusions of the WT versions of FL2 and ΔΕ2 plus their corresponding battery of mutated variants (see above and Fig.2) were expressed in HeLa cells and in *EPN2* K.O. mouse fibroblasts; next the transfectants were evaluated for the extent of proper colocalization with AP2 and mislocalization to the cytosol, TGN and nucleus. Once again, note that the latter is a well-known mislocalization artifact of free GFP (Fig.1B), this phenomenon is also observed when GFP is fused to proteins *lacking* strong localization determinants (*i.e.,* when GFP’s influence on localization becomes dominant) for example in the case of ENTH2-GFP (Fig.1B, 3^rd^ row).

Quantitative results obtained with ΔΕ and FL Epn2 mutant series are shown in Figs. 3 and 4, respectively. Representative images for the localization of these series can be found in Supplemental Figs.2 and 3 and Fig. 5. Importantly, although with lower levels of expression and colocalization, results obtained using *EPN2* K.O. fibroblasts (Fig. 5), validated observations made using the easier to culture and transfect HeLa cells; indicating that expression of endogenous Epn2 does not substantially alter the results obtained with the Epn2 mutated variants.

**Fig. 3.**
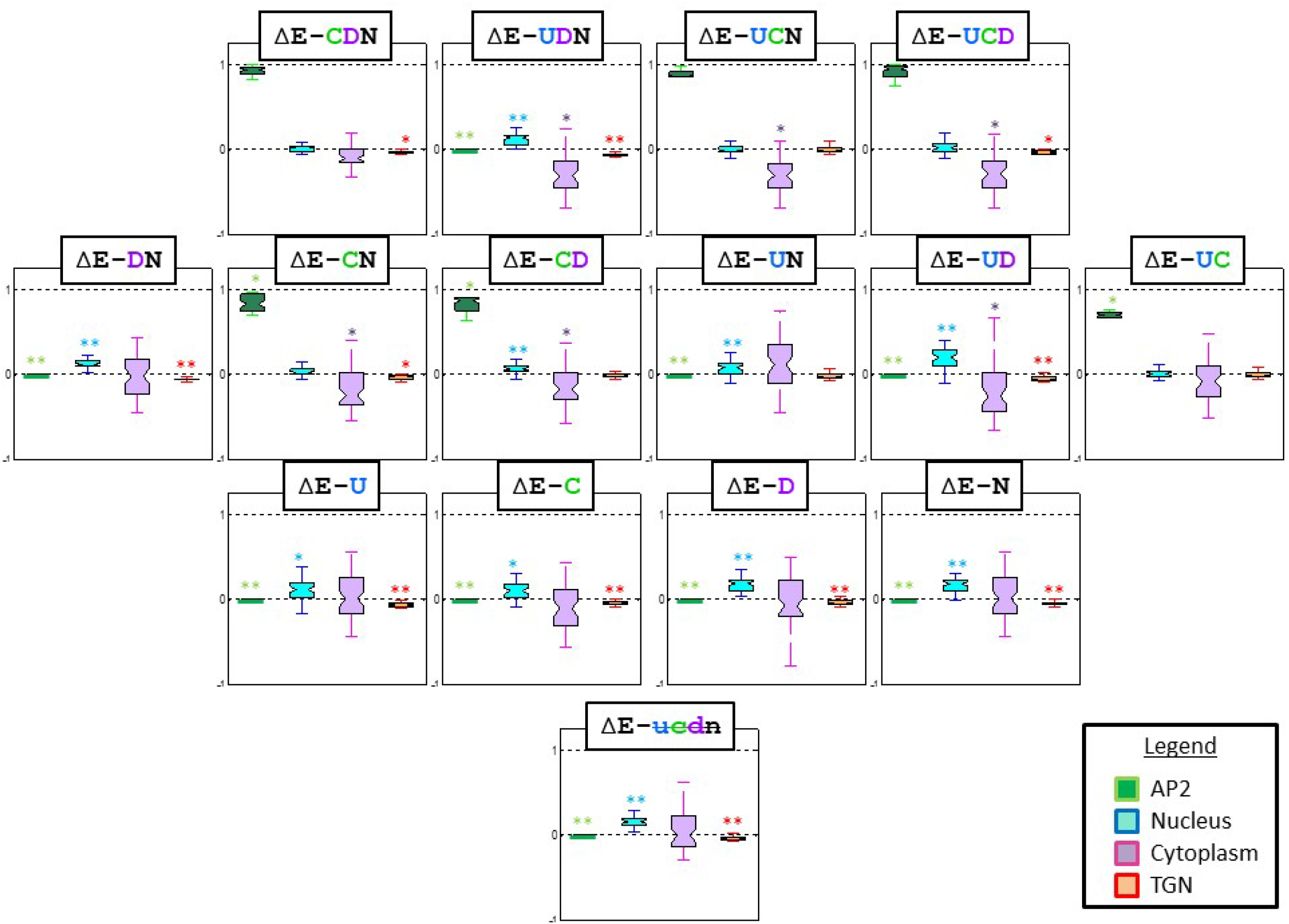
Quantitative results obtained for localization of the indicated ΔE variants of Epn2. Results for the indicated GFP-constructs (see Fig. 2) were obtained according to the procedure described in the *Results* and *Materials and Methods* sections. Please refer to text for details.

**Fig. 4.**
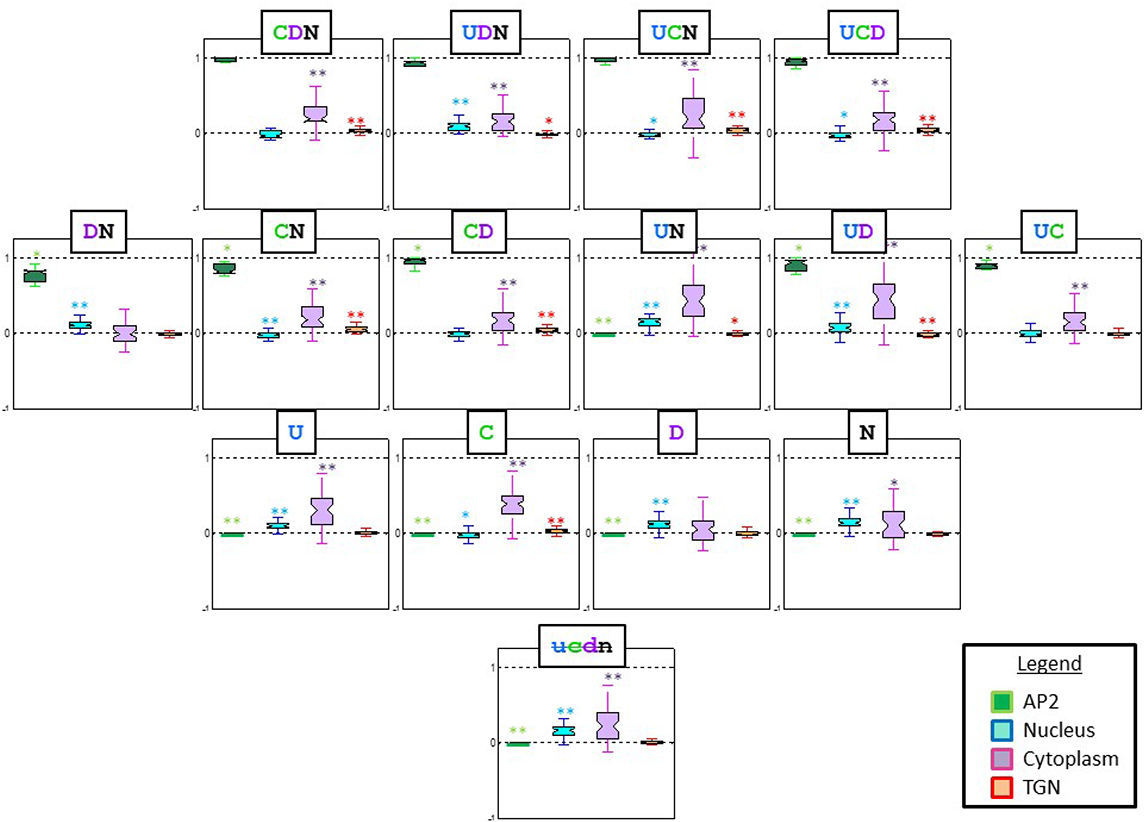
Quantitative results obtained for localization of the indicated FL variants of Epn2. Results for the indicated GFP-constructs (see Fig. 2) were obtained according to the procedure described in the *Results* and *Materials and Methods* sections. Please refer to text for details.

**Fig. 5.**
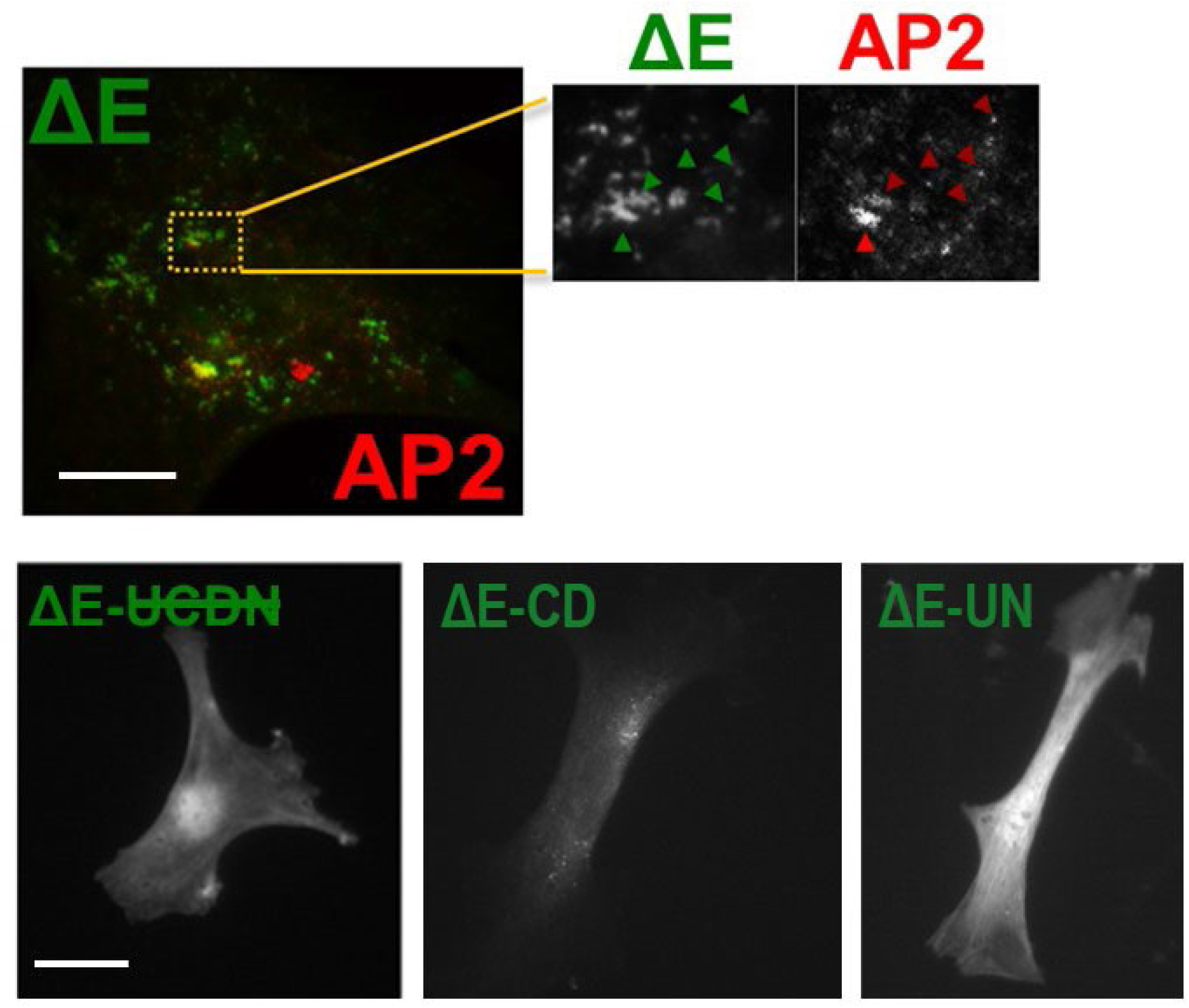
Localization of key ΔE Epn2 variants in *EPN2* KO cells. Skin fibroblasts isolated from *EPN2* knock-out mice were transfected for expression of the indicated ΔE Epn2 variants. Top row shows ΔE Epn2 colocalization with AP2. Box area is 3x enlarged on the right, arrowheads point to some examples of colocalization.

We first focused on the analysis of the results from the ΔΕ series:

Our results showed that in the absence of all 4 types of endocytic determinants (GFP-ΔE-2) the corresponding Epn2 mutated protein was unable to localize to AP2-positive endocytic sites in both HeLa and *EPN2* KO cells (Figs. 3 and 5 and Suppl. Fig.2). Indeed, virtually all protein partitioned between the cytosol and nuclear fractions, strongly resembling isolated GFP (Fig.1B); in other words, ΔΕ-2 showed only mislocalization (into the cytosol and dragged into the nucleus by GFP non-specific action). We concluded that although the short endocytic motifs account for <10% of the protein sequence they are absolutely required for proper localization to CME sites.

Concerning the intracellular distribution of the rest of the motif-mutant series, we observed the following: On the one hand, our data showed that presence of a single endocytic determinant (ΔΕ-U2, ΔΕ-C2, ΔΕ-D2 and ΔΕ-N2 variants) was not enough to support ΔΕ/AP2 colocalization (Fig. 3, Supplemental Fig.2B). However, we observed that ΔΕ-C2 (Epn2 truncation with only clathrin interacting motifs intact) exhibited a decreased nuclear fraction indicating a stronger ability to counteract the GFP tendency to translocate into this compartment.

On the other hand, half of the combinatorial mutants bearing two endocytic motifs were capable of at least partial colocalization to AP2-positive sites. Specifically, ΔΕ variants UC2, CD2 and CN2 showed substantial colocalization with AP2 (Figs. 3 and 5, Supplemental Fig.2B). The remaining three mutants in this group: ΔΕ-UD2, ΔΕ-UN2 and ΔΕ-DN2 were unable to target AP2-positive puncta (Figs. 3 and 5, Supplemental Fig.2B). The obvious difference between the AP2-colocalizing group and the ones that failed to target endocytic puncta is that in the former, one of the two available types of binding determinants were always the clathrin binding motifs (C). Strikingly, since the ΔΕ-UD2 and ΔΕ-DN2 variants failed at targeting AP2-positive endocytic sites, our results indicate that interaction *via* AP2 binding sites (DPW: D) was not sufficient for the proper localization of the Epn2 ΔE variants. These results are consistent with C being the stronger localization determinant. Mechanistic hypotheses suggested by our data are included in the *Discussion* section.

Next, we analyzed the mutants which had only one kind of endocytic determinant mutated, and therefore, the other three intact. All these mutants namely, ΔΕ-CDN2, ΔΕ-UDN2, ΔΕ-UCN2 and ΔΕ-UCD2 showed a substantial colocalization with AP2 (Fig.3 and Suppl. Fig.2), even in the absence of the C motifs (although a gain in nuclear signal was observed for the C-defective ΔΕ-UDN2 mutant indicating a weaker localization capability). These results indicate that although C is the strongest localization determinant category, other motifs within the carboxy-terminus of Epn2 can cooperate to target the protein to AP2-positive sites.

### The second C motif is more important than the first for Epn2 localization

Our results indicate that the clathrin binding motifs of Epn2 are its strongest localization determinants. However, it should be noted that the C-motifs in Epn2 fall into two different consensus classes, with the downstream 2^nd^ C-motif (C^b^=LVNLD) fitting a classical clathrin binding box **LФpФp** (where **Ф** represents hydrophobic residue and **p** represents polar residue) and the upstream one (1^st^ C or C^a^=LLDLMD) displaying a slightly different **LФpФФp** consensus (Drake et al., 2000). Interestingly, this array of C consensuses (a **LФpФФp** followed by a **LФpФp**) is conserved across mammalian Epns, opening the possibility of functional relevance. Therefore, to evaluate the role of each consensus and to test if this arrangement is required for functionality, we produce several additional ΔE mutated variants. Specifically, we capitalized the observation that while ΔE-UN2 was completely mislocalized, the ΔE-UCN2 (*i.e.,* with the “addition” of the C motifs) was highly efficiently targeted to endocytic sites (Figs. 3, 6 and suppl. Fig. 2). Therefore, we independently kept unaltered each individually C-motif (C^a^: LLDLMD and C^b^: LVNLD) in the otherwise mislocalized and C-deficient ΔE-UN2 construct and tested the new variants (bearing only one of the two C motifs: ΔE-UC^a^N2 and ΔE-UC^b^N2) for their ability to localize to AP2-positive structures. Our results showed that introduction of C^a^ yielded a 30% in endocytic site targeting, while presence of C^b^ produced a 70% colocalization with AP2 (Fig.6A), indicating that classical **LФpФp** consensus was more effective at inducing proper localization. Since in mammalian Epns the position of one and the other C consensus motif is also highly conserved (with C^a^ always immediately carboxy-terminal to the U motifs and the C^b^ just preceding the N determinants, see Fig. 1A), we tested if position of each C consensus motif has functional relevance. Specifically, we exchanged C-motif positions with and without 10-aminoacid flanking regions (*i.e.,* inserting C^a^ ± flanking region into the C^b^ position and vice versa) in the context of ΔE-UN2 (producing the ΔE-UC^a/B^N2 ΔE-UC^b/A^N2 variants, where a/B and b/A indicates C^a^ inserted into the native position of C^b^, and C^b^ inserted into the native position of C^a^, respectively) and proceeded to analyze their colocalization with AP2-positive structures. Strikingly, none of these constructs localized to AP2 endocytic sites (Fig.6B). Same results were obtained when C^a^ or C^b^ were introduced with their corresponding flanking regions (data not shown). These results indicate that both C^a^ and C^b^ motifs are functional only when at their native positions, perhaps due to conformational effects or cooperative interactions with other motifs or players. Although beyond the scope of this study, these hypotheses (among others) are currently being tested in our lab.

**Fig. 6.**
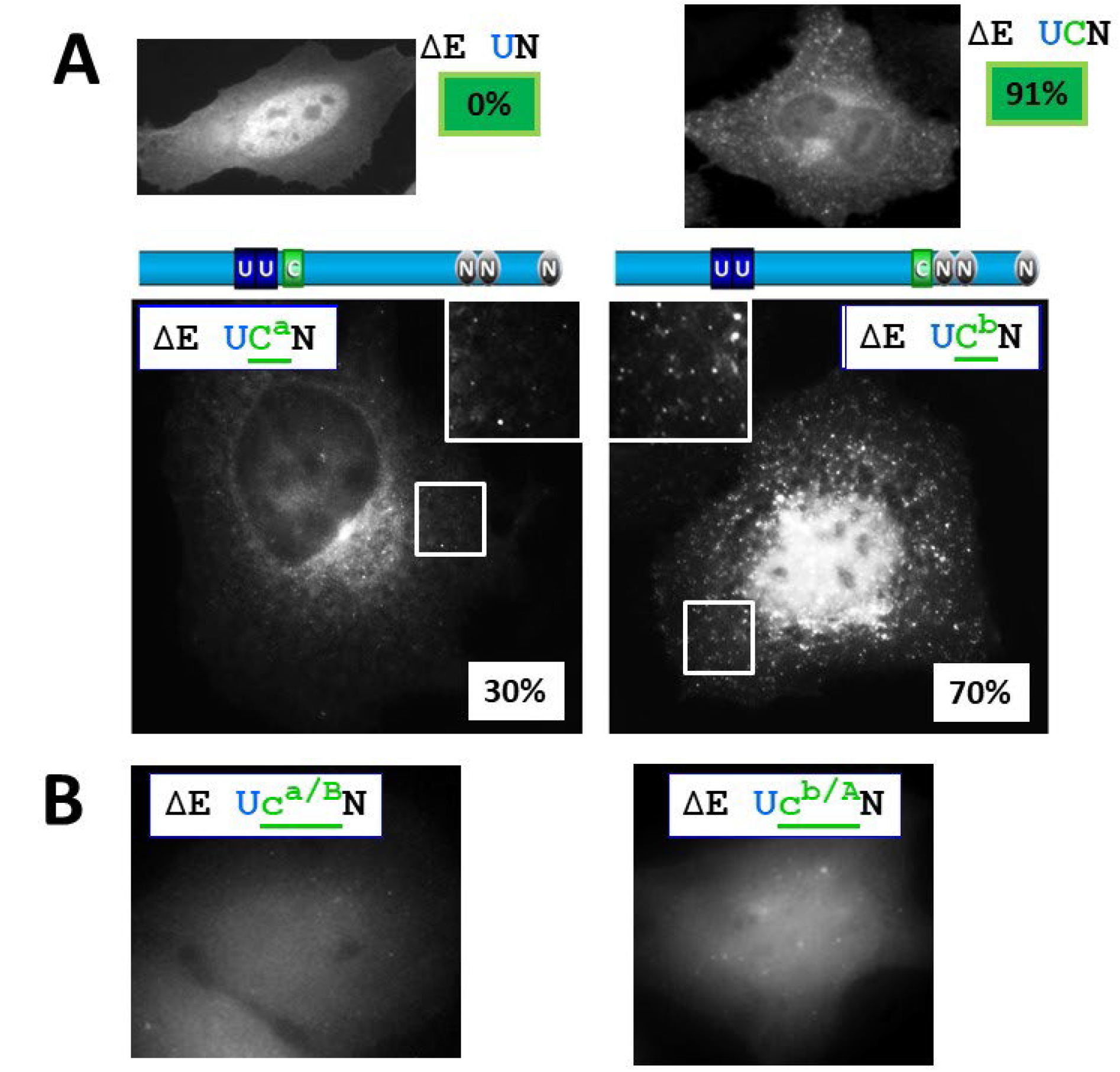
Relative relevance of the two C motifs of Epn2 for protein localization. **A**. Upper panels show the localization/AP2-colocalization capabilities of the ΔE UN vs UCN. Localization gain by presence of C was dissected by producing ΔE UC^a^N and ΔE UC^b^N variants (see diagrams included and text for more details), Representative images of both variants are shown along with quantitative results of colocalization with AP2. **B**. Introduction of C^a^ in the C^b^ position (UC^a/B^N) and viceversa (UC^b/A^N), resulted in no colocalization with AP2. See text for details.

### Establishment of a hierarchy for different endocytic determinants based on their relevance for Epn localization

The observation that three types (U, D and N) of endocytic binding motifs could cooperatively mediate Epn targeting to AP2-positive sites (see above) in the absence of clathrin-binding determinants (C), indicate that such motifs also contribute to some extent to protein localization. However, it was not possible for us to reliably assess the individual relevance of U, D and N motifs for Epn localization using ΔE constructs. A similar scenario was observed for the ENTH2 domain as it contributed to localization (FL2 was more efficiently localized than ΔE2, see above and Fig.1B) but was not a major player.

Therefore, we considered the ENTH2 domain as a 5^th^ localization determinant and proceeded to analyze whether the presence of the ENTH2 domain could also cooperate with U, D and N motifs to target endocytic sites. In other words, we compared the localization results obtained using ΔE vs FL Epn mutant series and tested the ability of the ENTH2 domain to enhance protein recruitment to AP2-positive sites (see Figs. 3, 4, and Suppl.Fig.2,3). Of particular importance were the results showing lack of endocytic site targeting observed with the C-lacking ΔE-UD2, ΔE-DN2 and ΔE-UN2 (see above) as compared to their counterparts generated using FL. Our results indicate that although ΔE-UD2 and ΔE-DN2 were unable to localize to AP2-positive puncta (even when AP2-binding, D motifs were present), FL-UD2 and FL-DN2 showed substantial targeting to endocytic sites (Figs. 3, 4 and Suppl. Figs.2,3). These results indicate that the ENTH2 contribution (in these FL mutated constructs) made possible proper localization. Although as shown in Figs. 3, 4 and Suppl.Figs.2,3, the ENTH domain on its own as ENTH2-GFP or in the 2-GFP construct could not secure localization at AP2 sites. Further, it should be noted that even when the ENTH2 domain was present, the FL-UN2 variant did not colocalize with AP2; indicating that the ENTH2 domain is not the only factor needed to make a mislocalized ΔE variant to target AP2 sites. Indeed, FL-UD2 and FL-DN2 (91% and 78% colocalization with AP2, respectively) differ from the mislocalized FL-UN2 in the fact that the latter lacks D motifs. Therefore, as a whole our results indicate that the AP2-binding, D motifs and the ENTH2 domain synergistically contribute to the protein localization. In addition, the relevance of U motifs seems to be slightly higher or equal than the one from N sequences (based on the extent of localization of UD2 vs DN2), yielding a hierarchy of Epn2 localization determinants: C> ENTH2-D>U≥N; indicating that Epn2 proper intracellular targeting heavily depends on clathrin recognition followed by a cooperative effect between D-mediated AP2 interaction and the PI(4,5)P_2_-binding ENTH2 domain.

### Similarities and differences for the use of localization determinants among Epn paralogs

Mammalian Epn paralogs present analogous domain/motif general architecture (Fig.1A) and they all require their carboxy-terminal region for localization at AP2-positive sites (only their ΔE constructs were able to colocalize with AP2, see Suppl.Fig.1).

Similar to Epn2, the endocytic binding motifs (U, C, D and N) were required for Epn1 and 3 localization, as their ΔE- variants were unable to colocalize with AP2 and showed a diffuse distribution pattern with partition of the protein between the cytosol and nucleus (Suppl.Figs.4,5).

We performed a systematic combinatorial mutagenesis of Epn1 and Epn3 followed by testing the resulting battery of mutated variants for colocalization with AP2 and mislocalization to cytosol/TGN/Nucleus (*i.e.*, analogously to the analysis described for Epn2), see Fig. 7 and Suppl. Figs.4,5.

**Fig. 7.**
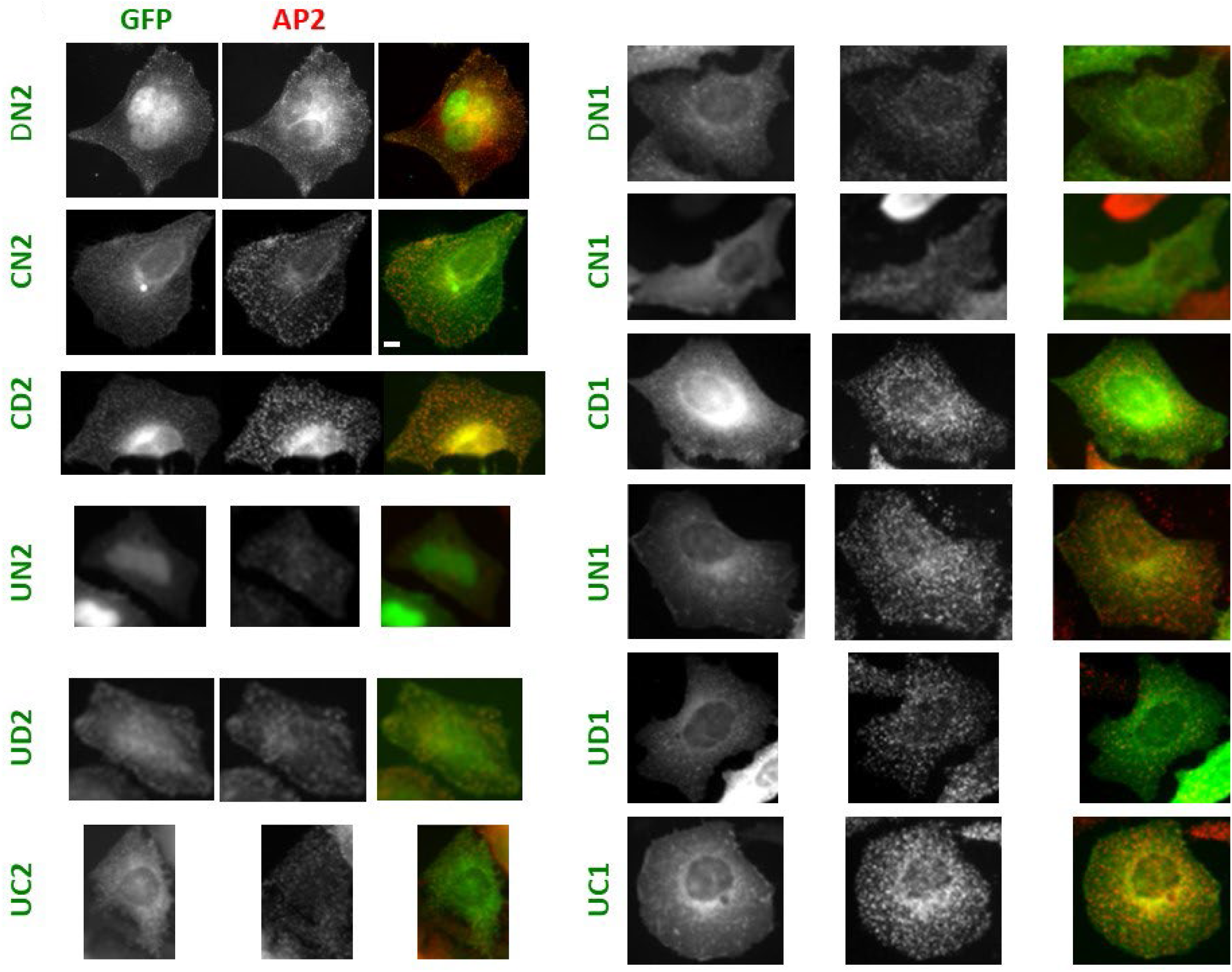
Comparison of the ability of FL Epn2 and Epn1 variants containing two endocytic determinants to colocalize with AP2. See text for details.

For Epn3, these studies led us to conclude that this paralog utilizes the same endocytic determinants for localization and with the same hierarchy than Epn2. Suppl. Fig.4 shows some critical Epn3 mutated variants that demonstrate such parallelism with Epn2. For example, out of the six variants having two types of endocytic determinants only the C-containing ΔE-CN3, -CD3 and -UC3, efficiently localized to AP2-positive sites (Suppl. Fig.4).

Epn1, 2 and 3 have paralog-specific regions (PSR) of uncertain function. However, these paralog-specific regions do not seem to substantially contribute to localization as their elimination did not affect the protein’s ability to target AP2-containing sites (data not shown).

The Epn1 paralog presented a slightly but important difference in motif hierarchy relevant to protein localization. Indeed, AP2 colocalization analysis showed that *none* of the GFP-ΔE1 constructs displaying two endocytic determinants (ΔE-DN1, ΔE-CN1, ΔE-CD1, ΔE-UN1, ΔE-UD1 and ΔE-UC1) colocalized with AP2 (in contrast to Epn2 and 3, where the C-containing motif variants were successfully targeted to CME sites), see Suppl. Fig.5, indicating that in contrast to Epn2/3, Epn1’s C motif is not the stronger localization determinants. Further, from all the GFP-FL1 corresponding mutants (FL-DN1, FL-CN1, FL-CD1, FL-UN1, FL-UD1 and FL-UC1) only the D motif containing proteins (FL-DN1, FL-CD1 and FL-UD1) were able to colocalize with AP2 (Fig. 7 and Suppl. Fig. 5). This results, once again highlight a synergism between the ENTH domain and the D motifs (as it was observed when analyzing Epn2 and Epn3 localization mutants).

Nevertheless, the main difference between the Epn paralogs is that for Epn1 the C-motifs represented weak localization determinants. Therefore, we speculate that left to rely mostly on the moderate ENTH-D motif synergism makes Epn1 the paralog with weaker localization determinants. Indeed, this may explain why FL-UCN1 even when displaying 3 different types of motifs (including C determinants) and the ENTH (but in absence of the ENTH-D motif synergism) cannot localize with AP2 endocytic sites. Nevertheless, we expect that due to avidity effects (Vauquelin and Charlton, 2013) the WT protein would have no problems to properly localize to sites of CME.

### Localization of Epn2 variants in endocytic pits versus plaques

Clathrin-containing endocytic structures can be categorized into two types, namely endocytic pits and endocytic plaques (Saffarian et al., 2009). Endocytic pits are smaller structures with shorter lifetimes of about 70sec seen in the dorsal part of the cell, while endocytic plaques are larger structures with longer lifetimes of ∼3mins on the face of the cell attached to the dish surface. On the one hand, the function of the pits is endocytosis of specific cargoes and constant membrane turnover in the leading edge of migratory cells (Kural et al., 2015, Saffarian et al., 2009). On the other hand, the presence of plaques is cell-dependent and is linked to cell-adhesion, where a loss of plaques resulted in an increase in cell migration (Saffarian et al., 2009, Kural et al., 2015).

Since Epn has been shown to be a part of both types of endocytic structures, we next asked if the different mutated variants can localize at pits and plaques and/or if they somehow affect the formation of such sites. Therefore, cells transfected with a selected group of Epn2 mutated variants were imaged on the dorsal versus the ventral side to quantify the density of the pits versus the plaques respectively (number of plaques or pits per unit area) using a spinning disk confocal microscope. As reported before by others, we observed that the dorsal and the ventral Hela cell surfaces were enriched for pits and plaques, respectively. The cells were also immunostained for AP2 to assess if the mutants affected the total number of CME sites per cell. The results from selected mutated variants indicated that the total number of endocytic sites (number of AP2 puncta either pits or plaques) per cell is not significantly affected by expression of Epn2 WT or mutated variant (Suppl. Fig.6A). In addition, our results indicate that the ability of the different Epn2 mutants to localize was similarly affected for pits versus the plaques (Suppl. Fig.6B).

### Epn mutants result in more abortive pits

Next we evaluated the impact of affected Epn localization on function. The effect of each Epn paralog and their corresponding battery of variants on the internalization of specific cargoes is difficult to directly measure due to the functional redundancy among the members of this adaptor protein family (*e.g.,* likely requiring a double or triple *EPN* K.O.; and even including *EPS15* K.O. for cargoes such as EGFR). In addition, the study of other Epn targets such as Notch ligands are further complicated by requiring specific two-cell juxtacrine systems for the physiological induction of endocytosis. Therefore, we instead focused on studying the impact of mutations on Epn’s role in endocytosis dynamics.

An endocytic pit can turn abortive, productive or persistent (Loerke et al., 2009), with defining different lifetimes of 5 to 30sec for abortive, ∼90 secs for productive and greater than 90sec for persistent pits. Epns, among others, have emerged as ‘checkpoint’ proteins involved in deciding the fate of endocytic sites (Mettlen et al., 2009b). Therefore, the inability of some variants to properly localize or to interact with other elements of the endocytic machinery could conceivably affect pit fate/maturation. Therefore, we studied the dynamics of the endocytic sites in HeLa cells expressing Epn2 WT and mutated variants using TIRF microscopy. Our results indicated that Epn mutants altered the ratio of productive to non-productive (abortive and persistent) sites (Fig.8A). Cells expressing Epn2 mutated variants presented a significant decrease in productive and an increase in non-productive sites as compared to the WT. Moreover, the differences with respect to WT were larger for the mutants bearing only two types of endocytic motifs versus variants with three kind of binding determinants (Fig.8A).

**Fig. 8.**
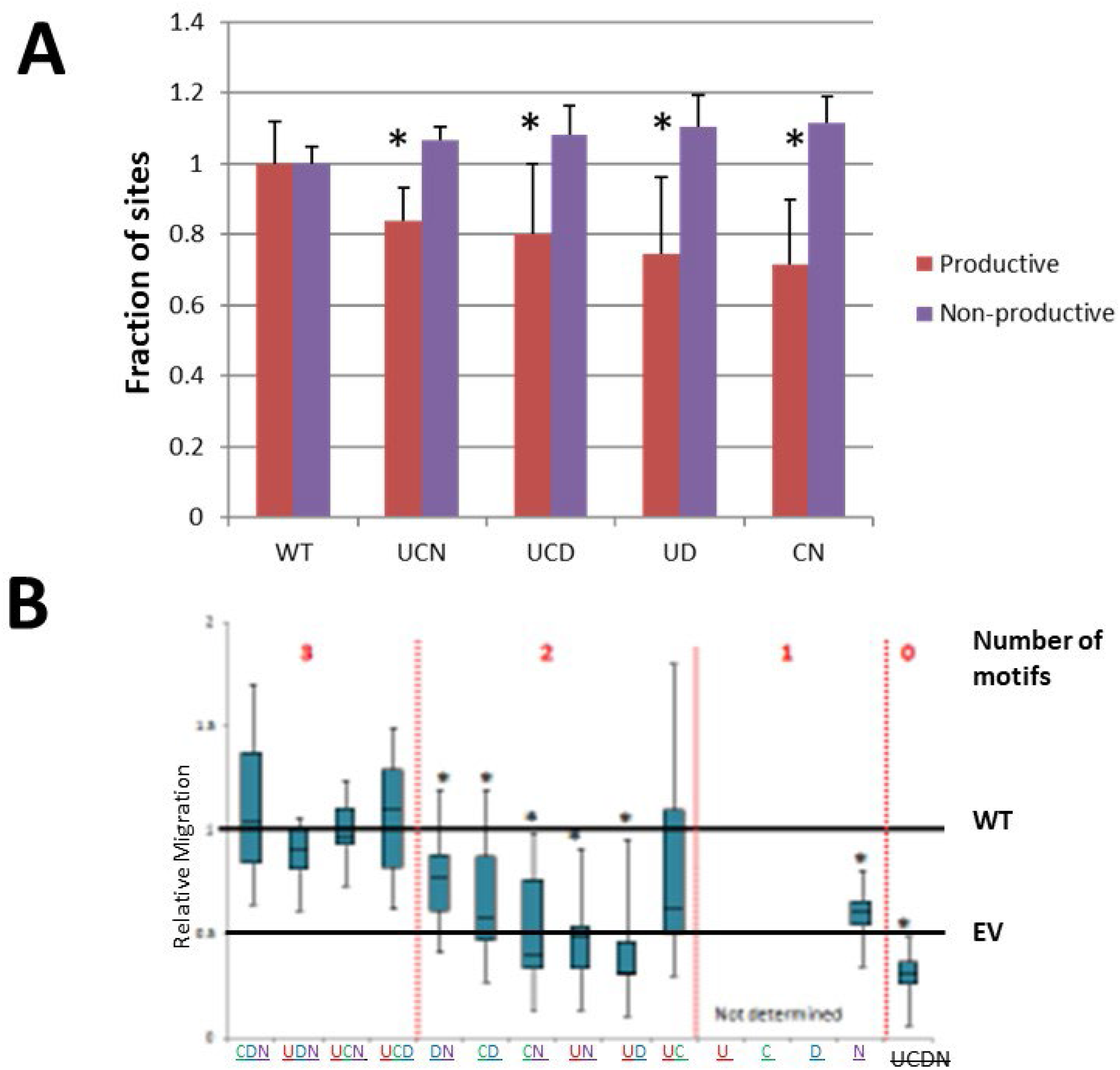
Functional relevance of different Epn2 endocytic determinants. **A.** Ability to sustain the assembly of productive endocytic pits. **B.** Ability to enhance cell migration. See text for details.

### Role of Epn carboxy-terminus in enhancing cell migration

We and others have established that upregulation of Epn2 and Epn3 expression enhances cell migration (Coon et al., 2010; 2011). Further, and although the presence of the ENTH domain is required, the carboxy-terminus is also needed.

Since we had established the role of the carboxy-terminus in Epn localization, we tested if a decrease in AP2 colocalization due to mutations in such region would affect the mutant’s ability to enhance cell migration. As mentioned before, the ENTH domain is required for enhancing migration; therefore, we expressed FL Epn2 WT and mutated variants and assessed the ability of the cells to migrate in transwells (Fig.8B). In general, we observed a correlation between proper localization of the Epn2 mutated variant and its ability to enhance cell migration (Fig. 8B); however, a few exceptions were noted. For example, although FL-CD2, -CN2 and -UD2 mutated variants had a substantial level of colocalization with AP2, they were impaired in their ability to enhance cell migration (Fig.8B). This observation also suggested that determinants like U and N, although not crucial for localization are required for proper function.

We also tested some representative Epn3 mutated variants to study the localization versus enhancement in migration correlation and observed a very similar trend to the one described for Epn2 (data not shown). The results suggest that Epn’s ability to enhance cell migration not only depends on the variant’s capability to target endocytic sites but also on interactions with other factors.

### *Saccharomyces cerevisiae’s* Epns relied on U and N motif for localization to endocytic sites

As indicated before, despite sequence and protein architecture differences (Fig.1A) the yeast Epns Ent1 and Ent2 (just like their human homologs) mainly relied on their unstructured carboxy-terminus for localization (Suppl. Fig.1). Nevertheless, it should be noted that there are several important differences between mammals and yeast concerning the process of endocytosis itself, *e.g*., *S. cerevisiae* shows less dependence on AP complexes (indeed, yeast Epns lack identifiable DPW/DPW-alike AP2-binding motifs) but also exhibits less reliance on clathrin and dynamin-like proteins, while requiring high involvement of the actin cytoskeleton for vesicle scission and showing differences in pit ultrastructural characteristics (Idrissi et al., 2008; Idrissi and Geli, 2014). Therefore, we investigated the overall use of Epn’s binding determinants for localization at yeast sites of endocytosis in comparison with their mammalian counterparts to evaluate if the differences mentioned above have parallel deviations in the mode by which these proteins are targeted to endocytic sites.

We took advantage of the availability of double Epn K.O. (also referred to as *Δent1, Δent2* or just *ΔΔ*—Aguilar et al., 2006) cells to test the ability of a series of Ent2-GFP mutants to localize at endocytic sites identified by presence of Abp1 (as yeast AP2 is not a valid marker for such structures).

Interestingly, our results indicate that a yeast Epn mutant lacking its only and very carboxy-terminal C motif (fitting a **LФpФp** consensus) was not impaired for proper localization in *ΔΔ* cells (UQN, Fig.9A). In fact, elimination of the region containing both N and C motifs had no substantial effect on the localization of the resulting GFP-Ent2 truncation (UQ). Further, as indicated before, the ENTH domain on its own (*i.e.,* a truncation lacking U, C, N motifs) was severely affected for targeting at endocytic sites; however, it was noticeable that in contrast to WT cells (Suppl. Fig.1), in *ΔΔ* cells a few scattered puncta were observed (Fig.9A, row 5), suggesting that the presence of endogenous yeast Epns easily competed off the recruitment of this weaker binder to Abp1 sites.

**Fig. 9.**
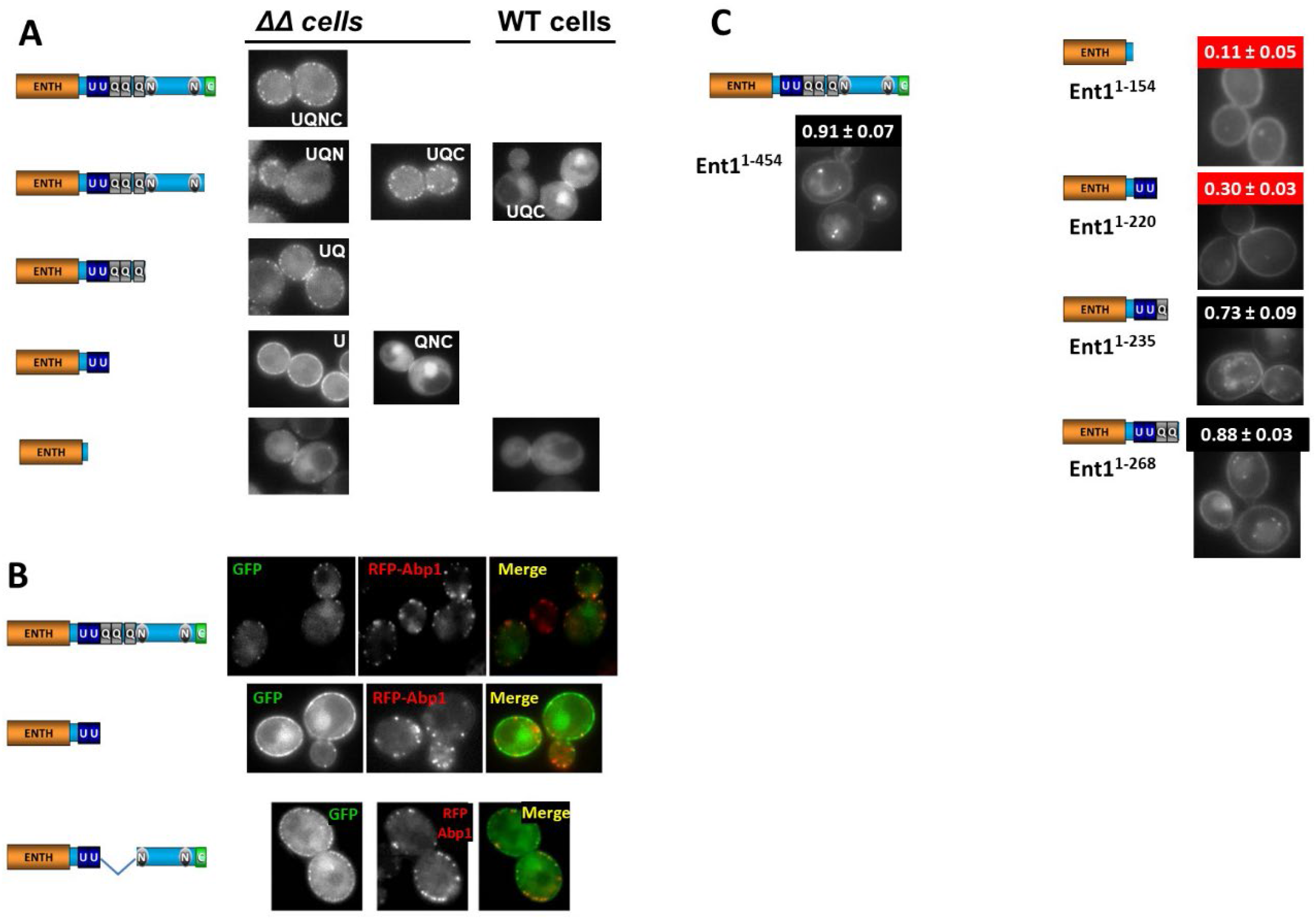
Role for localization and function of different determinants in yeast Epns. **A**. Localization of different GFP-Ent1truncations and point mutants in *ΔΔ* and WT cells. See text for details. **B**. Role of Q regions for GFP-Ent1 localization. See text for details. **C**. Relevance of Q regions for Ent1-mediated internalization of the Na^+^ pump Ena1. See text for details.

Since in *ΔΔ* cells C and N motifs were not required for localization, but a truncation lacking the U, C, N motifs (*i.e.,* the ENTH domain) could not target Abp1 structures, we hypothesized that U is important for recognition of endocytic sites; therefore, we tested a FL GFP-Ent2 variant with *only* its U motifs mutated (QNC). Importantly, our data indicate that this U-deficient Ent2 was highly deficient in its ability to localize at Abp1 sites (Fig.9A, row 4).

In addition, and although the C-deficient variant properly localized at Abp1 sites in both WT and *ΔΔ* cells (confirming that C motifs are not substantially contributing to yeast Epn localization), a FL variant with *only* the N motifs mutated (UQC) localized well in *ΔΔ* cells (supporting our previous observations with Ent2 truncations using the same strain—Fig.9A, row 2), but was affected for localization in WT cells (Fig.9A, row 2). This result indicated that the N motifs also participate in yeast Epn localization, but that they are very weak contributors which absence only affected recruitment to Abp1-structures upon competition exerted by the endogenous protein (Fig.9A, row 2).

Therefore, as a whole our results suggest a hierarchy of endocytic determinants for recruitment to endocytic sites: U>N>>C (*i.e.,* a virtually reverted order as compared to the one exhibited by human Epns).

### Role of yeast Epn’s glutamine-rich specific sequences in localization at endocytic sites

Importantly, we also observed that a U-containing, carboxy-terminal truncation lacking yeast homolog-specific glutamines rich (Q) sequences exhibited a unique, abnormal localization (Fig.9B row 2). Specifically, this truncation showed plasma membrane-localized discrete puncta but in a widespread pattern and more abundant than the Abp1-positive sites, *i.e.,* although showing a certain degree of colocalization with Abp1, a substantial amount of Ent2 variant-containing puncta was localized to structures containing no detectable Abp1 (and viceversa; Fig.9B).

To confirm that the effect was due to lack of Qs, we prepared another variant in which *only* the Qs were deleted (GFP-Ent2 ΔQ) and its colocalization with Abp1-sites was analyzed. We indeed observed that the GFP-Ent2 ΔQ variant also showed a pattern characterized by very abundant puncta and only partially colocalizing with Abp1 (Fig. 9B row 3). These results verified that lack of Qs caused the above-described abnormal localization phenotype, suggesting that that these sequences found in yeast Epns contribute to enhance specific targeting of plasma membrane Abp1 sites.

### Localization and function relationship for yeast Epns

We recently identified the Na^+^ pump Ena1 as the first yeast Epn-specific endocytic cargo, and we established that either Ent1 or Ent2 ΔE truncations (*i.e.,* the ENTH domain was not required for this function) could sustain the internalization of this membrane protein (Sen et al., 2020). Further, since Ena1’s uptake depended on recognition of its ubiquitinated K^1090^, integrity of Epn’s U motifs was an absolute requirement for Ena1 internalization (Sen et al., 2020). Given the findings described above, this observation is also likely to be due to defects in Epn localization to endocytic sites.

Here we expanded our studies to test the role of yeast Epn’s N motifs and Q regions in Ena1 Endocytosis. Specifically, we transformed *ΔΔ* cells expressing Ena1-GFP with plasmids for HA-tagged Ent1 variants (Suppl.Table II) and estimated the value of the so-called Intracellular Localization (IL) index.

To test the relevance of Qs for cargo internalization, we focused on Ent1 as it contains Q of shorter length than Ent2 (Fig. 1A) identified using a sequence analysis strategy developed in our lab (see Sen et al., IEEE and *Materials and Methods*) and prepared additional truncations. and proceeded to determine the Ena1 IL index in *ΔΔ* cells (Fig.9C). As expected, the presence of an ENTH domain-only variant did not increase the uptake of Ena1 above background levels; however, the addition of the U motifs tripled the Ena1 IL value (Fig.9C). As indicated before, the presence of the U determinant may increase in Ena1 internalization by improving the variant’s ability to properly localize at endocytic sites (Fig.9A) but also due to the gained capability of recognizing ubiquitinated-Ena1 (Sen et al., 2020). Nevertheless, it should be noted that Ena1 IL value obtained by expression of the ENTH-U (Ent1^1-220^) variant is substantially below the one observed in presence of FL Ent1 (Fig.9C). Importantly, addition of the first Q stretch (ENTH-UQ_1_: Ent1^1-235^) significantly increased (doubled) the IL of Ena1 in presence of ENTH-U (Fig.9C); further, the subsequent addition of the second Q region (ENTH-UQ_1_Q_2_ or just ENTH-UQ: Ent1^1-268^) brought the IL to values comparable to the ones measured under expression of FL Ent1 (Fig.9C). Indeed, shorter truncations of the carboxy-terminus also including the N and the C (Ent1^1-454^ FL) motifs, caused only moderate (non-statistically significant) increases in the uptake of Ena1over values obtained with ENTH-UQ. Ent2 was redundant with Ent1 for mediating Ena1 uptake (Sen et al., 2020) and exhibited a similar dependence on U motifs and homologous Qs for function (data not shown).

In summary, our results indicate that the presence of Q regions enhances the ability of yeast Epns to mediate Ena1 endocytosis.

## DISCUSSION

Biological function depends on proper temporal and spatial protein localization. Therefore, it is not surprising that the temporal sequence of protein recruitment at endocytic sites has been the focus of important work developed during recent years (Lu et al., 2016). Here we aimed to complement those and other studies by clarifying the process by which the Epn family of adaptor proteins targets sites of endocytosis. This knowledge would contribute to the understanding the mechanisms behind the essential process of endocytosis, and to interpret differences concerning such process in human vs yeast cells. This is particularly relevant as *S. cerevisiae* has proven to be a powerful genetic/cell biological system that has been and continue to be used as a model for higher eukaryotes. Nevertheless, it is now well-known that yeast presents important differences with other species (*e.g.,* humans) in terms of how cargo internalization takes place (REFs). Specifically, endocytosis in yeast is less dependent on clathrin, APs and dynamin-like proteins, but has a more intense use of the actin cytoskeleton than in mammals (REFs). Further, the ultrastructure of the clathrin-coated endocytic pit seems different in yeast. Indeed, constriction of the pit neck is less marked and the clathrin coat seems restricted to the base of the invagination (Fig.10). Therefore, an emerging question is: do proteins involved in endocytosis also differ in their mechanisms of recruitment to endocytic sites in yeast vs mammals?

**Fig. 10.**
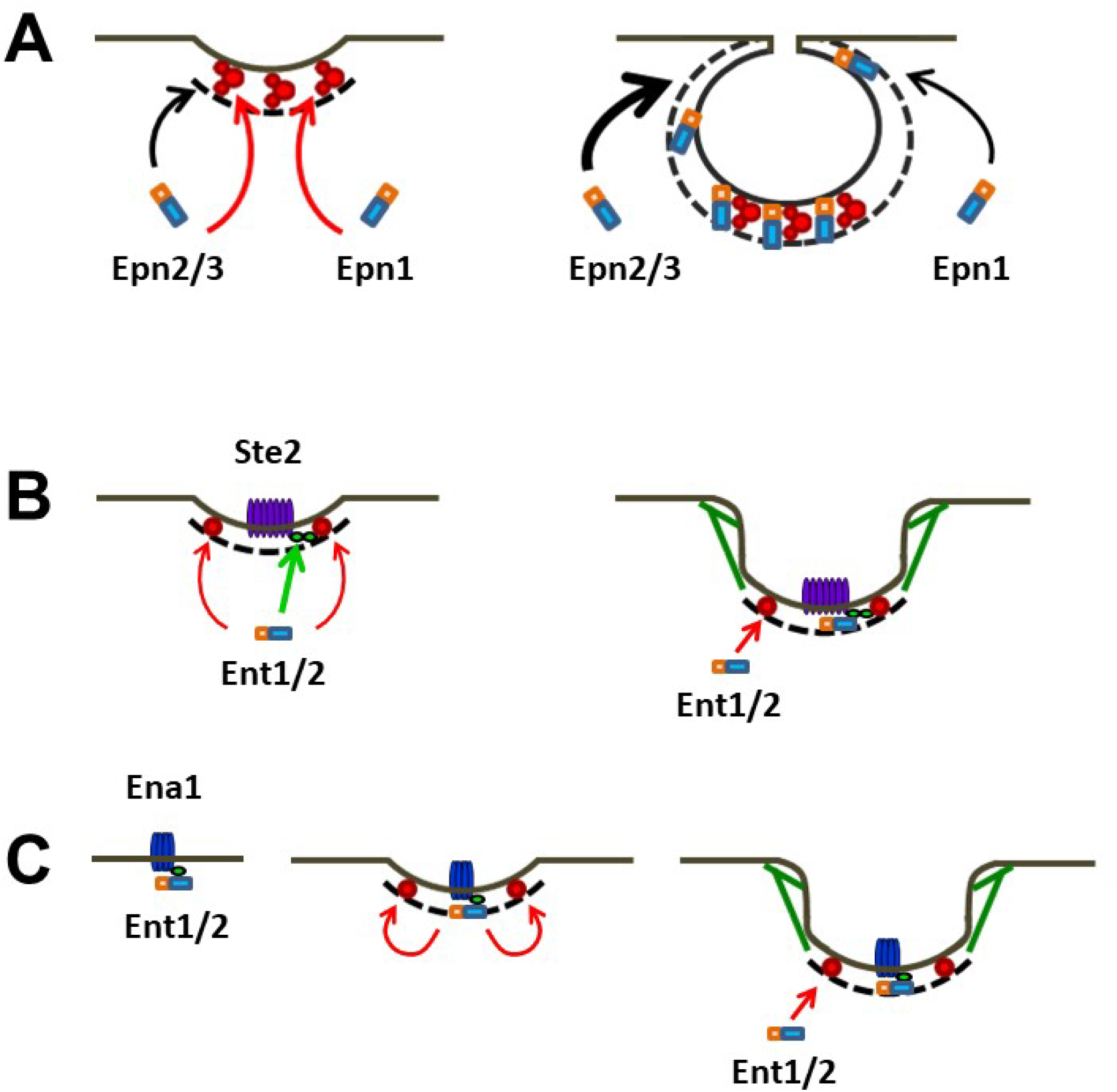
Working models for localization of human and yeast Epns at endocytic sites. **A.** *Left*: Epn2 and Epn3 target nascent endocytic sites by interacting with clathrin (black dashed line) and AP2 (red-colored complex); in contrast, Epn1 mainly relies on its ability to bind AP2. *Right*: At late maturation stages, additional Epn2/3 molecules can access the AP2-lacking upper segments of the endocytic pit *via* clathrin recognition, while recruitment of Epn1 is, in comparison, lesser. See text for details. **B-C.** Yeast Epns Ent1/2 localize to endocytic sites by engaging their U motifs with Ub (green ovals) covalently attached to cargoes. **B**. *Left*: Ent1/2 recognizes Ub chains conjugated to cargoes such as Ste2 (purple multispanning membrane protein) that can be retained in nascent endocytic sites by redundant non-Epn adaptors (*e.g.,* Ede1). Epn recruitment is further consolidated by N-mediated interactions with EH-containing early coat components such as Ede1 (red circles). *Right*: Epn recruitment at later stages of pit maturation is mostly mediated by interaction N motifs with EH-domain containing proteins Ede1 and Pan1 (red circles). **C**. *Left*: Ubiquitination of Epn-specific cargo Ena1 (blue multispanning membrane protein) allows Ent1/2 binding via U motifs. *Middle*: the resulting complex either initiate the assembly of a subgroup of endocytic sites or it associates with nascent clathrin pits *via* recognition of early component Ede1. *Right*: Epn recruitment at later stages of pit maturation is mostly mediated by interaction N motifs with EH-domain containing proteins Ede1 and Pan1.

Our studies shed light on the role for localization and function of different molecular determinants some which are common to yeast and mammalian Epns (*i.e.,* ENTH domain, U, C and N motifs) and others are homolog/paralog-specific (*e.g.,* D motifs and Q regions).

Overall, the results of the current work showed that:

1. The unstructured carboxy-terminus of Epn from both human and S. cerevisiae cells contain their major localization determinants.
2. While human Epn mostly relied on clathrin binding and/or an ENTH-D synergism for localization, the yeast homologs were more dependent on recognition of ubiquitinated cargo and EH domain-containing proteins to target endocytic sites.
3. Q sequences in the yeast homologs, although not sufficient on their own for protein localization, contribute to target endocytic sites.
4. Epn function requires proper localization but also the presence of specific determinants mediating necessary protein-protein interactions.

Below we discuss each of these general conclusions as well as more specific observations.

Finally, we propose temporal/spatial models for Epn function in human and yeast cells.

### 1- The unstructured carboxy-terminus of Epn from both human and yeast cells contain their major localization determinants

Our work indicated that a characteristic of all Epn paralogs and homologs is that their carboxy-termini contain the major determinants of localization with their ENTH domain playing a supporting role. On the one hand, although the human ENTH domain on its own showed some degree of interaction with the plasma membranes, we could not detect colocalization with AP2 in human cells. On the other hand, although the ENTH-lacking, mammalian ΔE truncation efficiently targeted endocytic sites, it also showed mislocalization at the TGN and such abnormal distribution was suppressed by mutation of the C motifs. These results suggest that in the absence of the PI(4,5)P_2_-binding ENTH domain, the ΔE variant has less selectivity for plasma membrane-vs TGN-localized clathrin. In addition, in mammals we detected cooperativity between the ENTH domain and D motifs for localization (see item 2 below).

One first obvious question arising from this work is: *since the ENTH domain binds to PI(4,5)P_2_ (which is enriched at endocytic sites), why is not capable of colocalizing with AP2?*

Our data obtained under *in-cell* physiological conditions, showed that the ENTH domain is highly cytosolic and is dragged by the GFP into the nucleus (as is to be expected for a protein with weak localization determinants). Indeed, this result was obtained for all ENTH domains from the different homologs and paralogs tested. This is consistent with studies conducted by others in living cells indicating that ENTH-GFP binds PI(4,5)P_2_ with low affinity (Leitner et al., 2019). We speculate that only when part of a multivalent and multispecific molecule (*e.g*., Epn^WT^ FL), the ENTH domain contributes to localization by enhancing avidity/matricity (Schmid et al., 2006). Further, an important functional consequence of this type of interaction is that the ENTH domain could participate as part of a multi-pronged coincidence detection system (see also item 2 below).

A second question posed by our data is: *why the residual interaction of the ENTH domain with the plasma membrane led to a widespread pattern?*

Addressing this question will require further dedicated investigations; however, we speculate that interactions of the ENTH domain with more widespread membrane components, *in the absence of coincidence detection*, causes this localization pattern. Whether these hypothetical components are lipids (for example other phosphoinositides, PIs) or proteinaceous components is uncertain. Interaction with other lipids might occur if the ENTH’s selectivity is relaxed enough (Aguilar et al., 2003); however, due to their transient and localized nature, PIs are not very abundant and most likely would still produce a plasma membrane discontinuous pattern. Perhaps, the involvement of some widely distributed proteins is more likely to be responsible for the ENTH membrane distribution. Although some interesting candidates exist (Coon et al., 2010), to identify the culprit(s) would require further dedicated investigation.

Importantly, the chief relevance of the carboxy-terminus for localization has also been observed in yeast. Indeed, a pioneering work from the Wendland lab (Maldonado-Baez et al., 2008) first reported such observation for Ent1, that here we confirmed and expanded.

Nevertheless, it should be noted that the Epn carboxy-termini from yeast and mammals differ in organization of their binding motifs (*e.g.,* one very carboxy-terminal in yeast vs two internal C motifs in mammals) and presence/absence of characteristic sequences (*e.g.,* D motifs and Q regions in mammals and yeast, respectively). Therefore, some differences in functionality can be expected (see items 2-4 below).

### 2- While human Epn relied on clathrin binding and an ENTH-D synergism for localization, the yeast homologs were more dependent on recognition of ubiquitinated cargo and EH domain-containing proteins to target endocytic sites

Human Epns required at least two functional motifs for recruitment to endocytic sites. Indeed, although C motifs are the strongest localization determinant for human Epn2/3, they were not capable of properly targeting the protein on their own and the presence of a second motif was needed. We observed that any other determinant (U, D or N) was able to complement C to promote localization, indicating that the effect likely results from the presence of additional binding motifs, strongly suggesting that induction of *avidity* by multivalent/multispecific molecules (due to kinetic, and other binding-enhancing effects—see Schmid et al., 2006; Vauquelin and Charlton, 2013) allowed specific membrane targeting. Further, the combination of all three U, D and N determinants together bypassed the absence of C (even in ΔE variants), also suggesting that combination of multiple different binding sites (providing multivalency/multispecificity) can induce localization likely due to avidity effects.

Importantly, a synergistic effect was noted specifically involving the ENTH domain and the D motifs in all human Epn paralogs. Although the nature of this cooperativity will also require further investigation, we speculate that results from coincidence detection of sites that contain PI(4,5)P_2_ and that also have a critical mass of AP2 in them.

Interestingly, the ENTH-D synergism is the main driver for the localization of the human Epn1 paralog in which the C motif (in contrast to Epn2/3) was not very influential. This is surprising because the C motifs themselves are quite similar between Epn1 and Epn2/3. Indeed, the three paralogs show the same LФpФФp(C^a^)—LФpФp(C^b^) consensus pattern and similar flanking regions. Further, it should be noted that at least for Epn2, we established that the LФpФp consensus (C^b^) was more impactful for localization, perhaps due to its higher affinity for clathrin (Drake and Traub, 2001). However, the less efficient C^a^ was evolutionary conserved at its upstream position; indeed, our results suggest that functionality requires the presence of both types of C consensuses in a specific configuration.

If C motifs are equivalent among Epn paralogs, it is possible that there are additional sequence elements that either downmodulate (in Epn1) and/or enhance (in Epn2/3) the effective binding to clathrin. Sequence analysis reveals candidate regions present in both Epn2 and 3 but absent in Epn1, for example a 12-residue stretch starting at position 174 and 185 in Epn3 and Epn2, respectively. However, the functionality of such sequence is uncertain, and this sequence is not immediately adjacent to the C motifs; therefore, if responsible for enhancing clathrin binding by Epn2/3, it would have to exert long distance intramolecular effects or being involved in recruiting a factor “X” which in turn would facilitate clathrin interaction with C motifs. Similarly, there is at least one sequence insert in Epn1 (starting at position 316; in between the two C motifs and within the D-containing region) not found in Epn2/3 that could, by an unclear mechanism, negatively affect clathrin binding. However, the insert is also no adjacent to any of the C motifs; therefore, also would require an at-distance intramolecular effect and/or the recruitment of a factor that somehow would downmodulate clathrin binding.

In summary, the experimental results are clear and compelling; however, the mechanism by which C motifs are downplayed (Epn1) and/or enhanced (Epn2/3) for targeting to endocytic sites, remains speculative. Nevertheless, it should be highlighted that, even when not critical, C motifs are expected to contribute to localization of the WT, FL Epn1 by adding to the molecule multivalency/multispecificity (*i.e., via* avidity effects).

In contrast to human Epns, the yeast homologs Ent1/2 did not rely on C, but mainly on U and on N motifs for targeting endocytic sites. We previously demonstrated that yeast Epn’s U motifs were required for internalization of the ubiquitinated (Ub)-Na^+^ pump Ena1 (Sen et al., 2020); therefore, they are expected to also play a role in cargo retention at nascent endocytic sites.

There are several hypothetical scenarios that combine both U motif roles (Epn localization and cargo binding); two *non-exclusive* possibilities are enunciated below:

a. *Ub-cargo is retained by non-Epn adaptors into nascent endocytic sites leading to Ent1/2 recruitment by Ub-U motif interactions.* This is consistent with the relatively late Epn recruitment observed by other investigators (Lu et al., 2016) and clearly applies to the case of cargoes such as Ub-Ste2 which is recognized by Ent1/2 but also by other adaptors such as Ede1 (Shih et al., 2002). This possibility obviously requires the availability of multiple Ub units in a single cargo molecule for simultaneous binding by different adaptors (other and Epns). The arrival of Epn to preassembled sites could be compatible with a maturation/checkpoint function for Ent1/2, similarly to what has been described in mammals. Further, N motifs could also engage in binding already present elements of the early coat (Ede1—Lu et al., 2016).
b. *Existence of a subset on endocytic sites in which yeast Epns play an earlier role*. This could be the case of Ub-Ena1 molecules which are recognized *only* by Ent1/2, *e.g.,* sites initiated by Epn recognition of cargo followed by incorporation to incipient stochastic endocytic patches (Ehrlich et al., 2004). This scenario would incorporate the roles of U motifs for the recruitment of cargo to endocytic sites and for localization at (and perhaps assembly of) a subset of Ena1-containing nascent endocytic sites.

Interestingly, human Epn1 isoform X contains not two (like all other isoforms/paralogs), but three U motifs; therefore, this feature predicts a more pronounced role for Ub-binding determinants in the localization of this variant while displaying its typical Epn1 C-independence. In other words, this specific Epn1 isoform is predicted to have localization characteristics similar to the ones showed from yeast Epns.

Finally, we did not observe ENTH cooperativity in yeast; however, a more detailed study would be required to rule out completely such possibility.

### 3- Q sequences in the yeast homologs, although not sufficient on their own for protein localization, they contribute to target endocytic sites

An interesting feature of several yeast endocytic proteins is the presence of stretches of sequence enriched in glutamine residues.

Polyglutamine regions have been implicated in protein-protein interactions; indeed, they have been shown to be able to form coiled-coil structures with “ambivalent hidrophobe” properties (*i.e.,* able to interact with both hydrophobic and hydrophilic residues). Therefore, we developed a procedure to identify regions within protein sequences with high propensity to form coiled-coils and to display Q-rich faces (Sen et al., 2017). Following such protocol, we defined 3 Q regions in both Ent1 and Ent2 (although length of the Qs were larger in Ent2; Fig. 1A). Remnants of such Q regions can also be found in Epns from other species (*e.g., D. melanogaster*), but not in human Epns.

Interestingly, our analysis (Sen et al., 2017) showed that in contrast to other Q-containing protein families (*e.g.,* transcription factors), yeast endocytic proteins frequently present Q regions also featuring non-Q residues (often acidic, *e.g.,* E) interspersed (Sen et al., 2017). In addition, we also observed that for endocytic proteins, at least the 2 initial turns of their coiled-coils also often display acidic residues that precede the Q-enriched sequence (Sen et al., 2017).

In fact, the 1^st^ Q region from both Ent1 and Ent2 is part of a single larger helical unit that starts with the 2^nd^ U motif including its terminal poly-E stretches (Fig. 1A). Therefore, considering that the U motifs are very important for yeast Epn localization, it is conceivable that integrity of the 1^st^ Q region could be important for U function or specificity. In agreement with this hypothesis, mutation of the pre-Q E-stretches affected protein association with the plasma membrane (Sen et al., 2017). In fact, protein localization was altered in a very similar manner to what it was observed upon elimination of the U motifs (Fig.9A, row 4), suggesting that E mutation causes U function impairment. Therefore, considering the results presented here and our previous work (Sen et al., 2017); we propose that the 1^st^ Q from Ent1/2 cooperates with the 2^nd^ U motif for mediating Epn localization.

Further, deletion of the whole Q regions themselves (leaving untouched the E-stretch between U and Q), caused a dramatic change in the specificity of yeast Epn recruitment. Therefore, we speculated that Q regions are also required for finding Abp1-positive structures, as in their absence they *also* localize to structures lacking such marker.

In addition, we speculated that these Abp1-negative structures may represent:

I. Early stages of endocytic site assembly that still lacks substantial actin polymerization.
II. previously undetected, and/or Epn-induced, Abp1-negative endocytic sites.

It should be noted that the existence of endocytic sites devoid of Abp1 has been described before; however, their nature and function remain unknown. Therefore, testing hypotheses I) and/or II) is the focus of intense research in our lab.

### 4- Epn function requires proper localization but also specific determinants likely to be mediating necessary protein-protein interactions

Previous studies in mammalian cells have shown that endocytic sites are initiated by random sampling the membrane by stochastic formation of clathrin/AP2 patches (Ehrlich et al., 2004). In addition, recruitment of Epn to endocytic sites has been shown to shadow the localization of clathrin and AP2 at endocytic plaque and pit sites (Saffarian et al., 2009). In fact, our results, at least for Epn2, indicate that motif mutations have a similar impact on protein localization to both plaques and pits (Suppl. Fig.6). Proper localization is a pre-requisite for function; however, we also found that some Epn determinants are independently required for functions such as enhancement of cell migration and endocytic site maturation. Indeed, even though some mutated variants could localize to endocytic sites with high efficiency, they were unable to enhance cell migration and to contribute to endocytic site maturation. These results suggest that in mammals Epn function require proper localization (and likely endocytic function), but also that some determinants are important for other function exerted by Epns.

Concerning *S. cerevisiae*’s Epns: on the one hand, our results suggest a correspondence between localization and function; specifically, the U motifs are key for both. Indeed, lack of functional U leads to mislocalization of Ent1/2 and to absence of internalization of the yeast epn-specific cargo Ena1. On the other hand, the poorly localized, isolated ENTH domain is sufficient to sustain viability in the Epn double K.O. cells (Aguilar et al., 2006). However, the latter might be compatible with a scenario in which a small (almost undetectable) fraction of properly localized ENTH is enough to exert its endocytosis-independent function required for cell viability.

### Working models

Based on our results and the work of others we propose the following working models for Epn localization and function in human and yeast cells, at early and late stages of endocytic site formation:

**In mammals** we envision that **at early phases of endocytic site formation**, Epns targeting occurs *via* clathrin and AP2 binding (the latter synergistically enhanced by presence of the ENTH domain) to assure ub-cargo retention and endocytic network stabilization (Fig.10A, *left*).

Therefore, all Epn paralogs are expected to be homogenously distributed in the nascent clathrin-coated area (Fig. 10A). This prediction agrees with the observed localization of Epn in the AP2 cap of the developing endocytic pit.

**At later stages**, since AP2 is likely concentrated in the distal vesicle pole (Fig. 10A, right— Saffarian and Kirchhausen, 2008; Cheng et al., 2007), Epn recruitment at the upper, neck-proximal part of the pit is mostly driven by interactions with clathrin (Fig. 10A, *right*) where it contributes to membrane bending preceding vesicle scission.

In the case of Epn2/3, preference for C-over D-mediated interactions may assure robust recruitment all throughout pit maturation stages and to the different regions of the endocytic structure. At first sight, this seems to predict that Epn1 would have a minor role at the pit’s neck region. However, and despite the less prominent role of C motifs, avidity-driven recruitment would still take place via C and N motifs for the FL, WT Epn1. Interestingly, it has been suggested that clathrin binding impairs Ub-cargo interaction with Epn (Chen and De Camilli, 2005); therefore, Epn function at the upper part of the pit might be restricted to membrane bending and network stabilization. Of course, independently of when and how Epns were recruited to the pit, these adaptors are expected to contribute to maintain a functional maturation checkpoint (REF).

**In *S. cerevisiae***; however, we speculate (as discussed under item #2) that interaction with Ub-cargo and early coat components like the EH domain-containing protein Ede1 play a more important role in recruitment (Fig. 10C). As also mentioned before (item 2 above) we speculate on the existence of at least two non-mutually exclusive scenarios: a) yeast Epns are recruited *via* their U motifs into sites of endocytosis by Ub-cargo retained by redundant non-Epn adaptors (e.g., Ub-Ste2—Shih et al., 2002) followed by N-mediated interactions. b) Recognition of Epn-specific Ub-cargo (*e.g.,* Ub-Ena1; Sen et al., 2020) and initiation of endocytic site assembly or function.

In summary, although yeast and human Epns display similar endocytic protein binding motifs, they exhibit different relevance for proper protein localization and function. We believe that these differences run parallel to mechanistic divergences in the process of endocytosis in these organisms. As expected, out of these findings more questions emerge in terms of mechanistic details in one and the other organism. Current and future investigations in our and other labs will address such gaps ultimately yielding a more complete understanding of the localization-function relationship for proteins involve in endocytosis.

## MATERIALS AND METHODS

**Reagents:** Materials were purchased from Fisher Scientific (Fairlawn, NJ) or Sigma (St. Louis, MO) unless stated otherwise.

### Cells utilized in this study, culture conditions and transfection/transformation procedures

Hela cells were cultured in DMEM, Streptomycin/Penicillin, 2mM L-Glutamine and 10% fetal bovine serum (FBS) at 37°C in a 5% CO_2_ incubator.

*EPN2 Knock Out (K.O.) fibroblasts:* We isolated skin fibroblasts from EPN2 K.O. animals (Chen et al., 2009) as described before (Dell’Angelica et al., 2000). Briefly, animals were sacrificed by cervical dislocation, soaked in EtOH 70% and placed on a sterile, clean surface. Several pieces of endodermis were isolated from the abdominal area using sterile scissor and forceps and collected in a plastic 10cm plate containing DMEM, Streptomycin/Penicillin, 2mM L-Glutamine and 20% FBS. Holding the pieces against the bottom of the plate with a sterile forceps, the tissue was minced using a sterile scalpel until 2-3mm pieces were obtained while carving grooves on the plastic. Samples were kept on a 37°C, 5% CO_2_ incubator changing media periodically. After 4-8 days fibroblasts started to emerge from the grooves, once a good number of cells was observed, they were trypsinized, expanded and frozen stocks were prepared.

#### Yeast culture conditions and transformation procedures

laboratory W303 (MATα *ade2-1 his3-1 leu2-3112 trp1-1 ura3-1 can1-100*) and *ent1Δent2Δ* (MATa *ent1::HIS3 ent2::HIS3 leu2-3,112 ura3-52 his3-200 trp1-901 lys2-801 suc2-9*) strains were grown overnight at 30°C with shaking at 250 RPM in standard yeast extract–peptone–dextrose (YPD) or synthetic selective (amino acid dropout) media supplemented with dextrose and lacking appropriate amino acids for plasmid maintenance. Yeast was transformed by the Li-Acetate method following standard techniques.

#### DNA Manipulations/Mutagenesis

Plasmids used in this study are listed in supplemental tables I and II. DNA constructs were prepared using standard techniques. Site directed mutagenesis was done using a QuikChange^®^ Lightning Site Directed Mutagenesis kit (Agilent Technologies, Inc., Santa Clara, CA).

### Quantification of colocalization between GFP-tagged Epn variants and AP2

Random images totaling approximately 200 punctate structures in at least 5 cells were analyzed for each variant and were repeated at least twice. Cells having only one source of fluorescence (expressing a GFP fusion but not immunostained for AP2, and viceversa) were used to estimate the amount of signal detected in channel “A” that corresponds to non-filtered fluorescence from channel “B” (crosstalk). Graphs representing crosstalk intensity in channel “A” vs. fluorescence intensity in channel “B” were constructed, after linear regression, thresholds were set up at values T_A_ = (M+SD_M_)xF_B_ + (I+SD_I_) (where T_A_: threshold applied to channel A; M and SD_M_: estimates of mean and standard deviation of the slope; F_B_: total fluorescence intensity in the channel B; I and SD_I_: estimates of mean and standard deviation of the intercept to the origin). Fluorescence signal in co-stained cells with intensity above T_A_ were considered to be specific. Discrete structures with fluorescence intensity values above the cytoplasmic fluorescence intensity in Channel “A” were considered as puncta/organelles. The percentage colocalization was calculated using the formula below,

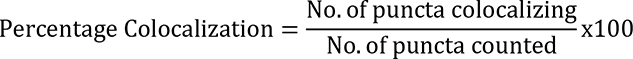

To measure nuclear and TGN localization, we stained the cells with DAPI or immunostained with antibodies against the TGN marker TGN46, respectively, and imaged along with the mutant (GFP-tagged) to be studied. Tracing the nucleus in the DAPI or the TGN channels and measuring the fluorescence intensity in the GFP channel within such region of interest produced the Epn-associated, nuclear or TGN fluoresce intensities. Tracing the perimeter of the cell and measuring the intensity in the GFP channel gave the total cell intensity. Tracing four cell regions devoid of puncta (with identical area), obtaining the mean fluorescence intensity value and integrating over the whole cell area gave the total cytoplasmic fluorescence intensity.

We analyzed 4 components for each variant under study: percentage of puncta colocalizing with AP2, percentage of total fluorescence intensity localizing in the nucleus, in the TGN and in the cytoplasm. Note that percentage of colocalization is not additive with the percentages deriving from the fraction of signal found in the nucleus, TGN and cytoplasm. Evaluation of the fraction of fluorescence associated with puncta is unreliable due to saturation effects and limitations in the estimation of individual and collective puncta size. Given that these measurements did not follow normal distributions, values were represented as independent boxplots for every sample. Puncta and region tracing was done manually using the Image J software.

#### Microscopy of yeast cells

Images were acquired using a Zeiss Axiovert 200M microscope equipped with Zeiss Axiocam MRm monochrome digital camera and Carl Zeiss Axiovision image acquisition software. For imaging *MET25* promoter driven constructs, cells were grown overnight in selective media in the presence of 2mM methionine. The cultures were diluted, washed with sterile H_2_O and protein expression was induced by growing the cells in selective media without methionine for 5h. A cell culture volume containing the equivalent of 2-5 OD_600nm_ of cells expressing fluorescently-tagged proteins were pelleted and resuspended in 50-100μl selective media, 10μl of cell suspension was spotted on a pre-cleaned slide, covered with a 22X22mm coverslip and imaged with appropriate filters as ∼0.3μm-spaced Z-stacks. Colocalization of GFP-Epn variant with Abp1-RFP was performed as described above.

### Dynamics of endocytic sites

Cells were transfected with GFP-tagged Epn WT or variants and seeded in live cell imaging chambers 12h before the experiment. After attaching, they were starved for 4h. Just before imaging, they were stimulated with serum containing media and sealed with parafilm. Imaging was done using a spinning disc confocal microscopy for 30min. Each cell was imaged for 3min on the dorsal surface at an interval of 1sec. 6 movies per sample were obtained. The movies were analyzed using the Image J plugin Speckle Tracker J (Smith et al., 2011). Only pits that were initiated after frame 1 and which disappeared before the last frame were considered for the analysis. 500-1000 sites per mutant were analyzed.

### Migration Assay

In this assay, 10^4^ mammalian cells were trypsinized, washed, resuspended in 500µl media lacking FBS and allowed to recover in suspension for 1h. Then cells were applied on upper chamber of 0.33cm^2^, 8µm pore transwell inserts (Corning Inc) coated with 100µg/ml BSA and 10µg/ml fibronectin on the upper and bottom side of the insert membrane, respectively. The migration chambers were then placed in wells containing media with 10% FBS in the lower chamber and allowed to migrate for 3h and fixed in 3% formaldehyde for 10min. Cells on fixed membranes were then stained with DAPI and visualized by epifluorescence microscopy. The number of transfected cells per membrane was counted in order to obtain cell inputs, and then the upper side of the membrane was wiped with a cotton swab and rinsed with PBS. Cells on the bottom side of the membrane were then counted and scored as migratory if their nuclei passed through the membrane pores. Percentage migration was determined as the ratio of migrated cells to the total number of transfected cells. Migration results were normalized based on WT percentage migration and expressed as mean ± SD of measurements from at least triplicate inserts.

#### Statistical analysis

The analysis was performed as described as described below and in (Taylor, 1997). When appropriate, the magnitude of errors associated with values derived from algebraic operations using experimentally measured quantities were calculated following standard rules of error propagation. The student’s t-test was used to evaluate differences among normally distributed ILs, while the Wilcoxon’s test was used to evaluate the significance between non-normal P/T of value samples. Bonferroni’s correction for multiple comparisons was performed whenever applicable [αC=p/n; n being the number of comparisons].

---

**SUPPLEMENTAL TABLE I:** PLASMIDS USED IN THIS STUDY

| **Plasmid** | **Description** | **Source** |
| --- | --- | --- |
| GFP | pEGFP-C1 | Clontech |
| FL1-GFP | Epn1 in pEGFP-N1 | H.McMahon (MRC Cambridge) |
| FL2-GFP | Epn2a in pEGFP-C1 | P. De Camilli (Yale) |
| FL2-mCherry | Epn2 in mCherry-C1 | Taylor et al, 2011 |
| FL3-GFP | Epn3 in pEGFP-C1 | B.Pilcher (Bioform Medical) |
| ENTH1-GFP | Epn1<sup>1-168</sup> pEGFP | Coon et al., 2010 |
| ENTH2-GFP | Epn2<sup>1-160</sup> pEGFP | This study |
| ΔE1-GFP | Epn1<sup>166-550</sup> pEGFP | Coon et al., 2010 |
| ΔE2-GFP | Epn2<sup>160-640</sup> pEGFP | This study |
| ΔE3-GFP | Epn3<sup>176-633</sup> pEGFP | This study |
| CDN1-GFP | Epn1 U mutated in pEGFP | This study |
| UDN1-GFP | Epn1 C mutated in pEGFP | This study |
| UCN1-GFP | Epn1 D mutated in pEGFP | This study |
| UCD1-GFP | Epn1 N mutated in pEGFP | This study |
| DN1-GFP | Epn1 U and C mutated in pEGFP | This study |
| CN1-GFP | Epn1 U and D mutated in pEGFP | This study |
| CD1-GFP | Epn1 U and N mutated in pEGFP | This study |
| UN1-GFP | Epn1 C and D mutated in pEGFP | This study |
| UD1-GFP | Epn1 C and N mutated in pEGFP | This study |
| UC1-GFP | Epn1 D and N mutated in pEGFP | This study |
| U1-GFP | Epn1 C, D and N mutated in pEGFP | This study |
| C1-GFP | Epn1 U, D and N mutated in pEGFP | This study |
| D1-GFP | Epn1 U, C and N mutated in pEGFP | This study |
| N1-GFP | Epn1 U, C and D mutated in pEGFP | This study |
| UCDN1-GFP | Epn1 U, C, D and N mutated in pEGFP | This study |
| ΔE-CDN1-GFP | Epn1<sup>166-550</sup>U mutated in pEGFP | This study |
| ΔE-UDN1-GFP | Epn1<sup>166-550</sup>C mutated in pEGFP | This study |
| ΔE-UCN1-GFP | D mutated in pEGFP | This study |
| ΔE-UCD1-GFP | Epn1<sup>166-550</sup>N mutated in pEGFP | This study |
| ΔE-DN1-GFP | Epn1<sup>166-550</sup>U and C mutated in pEGFP | This study |
| ΔE-CN1-GFP | Epn1<sup>166-550</sup>U and D mutated in pEGFP | This study |
| ΔE-CD1-GFP | Epn1<sup>166-550</sup>U and N mutated in pEGFP | This study |
| ΔE-UN1-GFP | Epn1<sup>166-550</sup>C and D mutated in pEGFP | This study |
| ΔE-UD1-GFP | Epn1<sup>166-550</sup>C and N mutated in pEGFP | This study |
| ΔE-UC1-GFP | Epn1<sup>166-550</sup>D and N mutated in pEGFP | This study |
| ΔE-U1-GFP | Epn1<sup>166-550</sup>C, D and N mutated in pEGFP | This study |
| ΔE-C1-GFP | Epn1<sup>166-550</sup>U, D and N mutated in pEGFP | This study |
| ΔE-D1-GFP | Epn1<sup>166-550</sup>U, C and N mutated in pEGFP | This study |
| ΔE-N1-GFP | Epn1<sup>166-550</sup>U, C and D mutated in pEGFP | This study |
| ΔE-UCDN1-GFP | Epn1<sup>166-550</sup>U, C, D and N mutated in pEGFP | This study |
| CDN2-GFP | Epn2 U mutated in pEGFP | This study |
| UDN2-GFP | Epn2 C mutated in pEGFP | This study |
| UCN2-GFP | Epn2 D mutated in pEGFP | This study |
| UCD2-GFP | Epn2 N mutated in pEGFP | This study |
| DN2-GFP | Epn2 U and C mutated in pEGFP | This study |
| CN2-GFP | Epn2 U and D mutated in pEGFP | This study |
| CD2-GFP | Epn2 U and N mutated in pEGFP | This study |
| UN2-GFP | Epn2 C and D mutated in pEGFP | This study |
| UD2-GFP | Epn2 C and N mutated in pEGFP | This study |
| UC2-GFP | Epn2 D and N mutated in pEGFP | This study |
| U2-GFP | Epn2 C, D and N mutated in pEGFP | This study |
| C2-GFP | Epn2 U, D and N mutated in pEGFP | This study |
| D2-GFP | Epn2 U, C and N mutated in pEGFP | This study |
| N2-GFP | Epn2 U, C and D mutated in pEGFP | This study |
| UCDN2-GFP | Epn2 U, C, D and N mutated in pEGFP | This study |
| ΔE-CDN2-GFP | Epn2<sup>160-640</sup>U mutated in pEGFP | This study |
| ΔE-UDN2-GFP | Epn2<sup>160-640</sup>C mutated in pEGFP | This study |
| ΔE-UCN2-GFP | Epn2<sup>160-640</sup>D mutated in pEGFP | This study |
| ΔE-UCD2-GFP | Epn2<sup>160-640</sup>N mutated in pEGFP | This study |
| ΔE-DN2-GFP | Epn2<sup>160-640</sup>U and C mutated in pEGFP | This study |
| ΔE-CN2-GFP | Epn2<sup>160-640</sup>U and D mutated in pEGFP | This study |
| ΔE-CD2-GFP | Epn2<sup>160-640</sup>U and N mutated in pEGFP | This study |
| ΔE-UN2-GFP | Epn2<sup>160-640</sup>C and D mutated in pEGFP | This study |
| ΔE-UD2-GFP | Epn2<sup>160-640</sup>C and N mutated in pEGFP | This study |
| ΔE-UC2-GFP | Epn2<sup>160-640</sup>D and N mutated in pEGFP | This study |
| ΔE-U2-GFP | Epn2<sup>160-640</sup>C, D and N mutated in pEGFP | This study |
| ΔE-C2-GFP | Epn2<sup>160-640</sup>U, D and N mutated in pEGFP | This study |
| ΔE-D2-GFP | Epn2<sup>160-640</sup>U, C and N mutated in pEGFP | This study |
| ΔE-N2-GFP | Epn2<sup>160-640</sup>U, C and D mutated in pEGFP | This study |
| $\Delta E$-UCDN2-GFP | Epn2<sup>160-640</sup>U, C, D and N mutated in pEGFP | This study |
| $\Delta E$-UC1N2-GFP | Epn2<sup>160-640</sup>D and C2 mutated in pEGFP | This study |
| $\Delta E$-UC<sup>b</sup>N2-GFP | Epn2<sup>160-640</sup> C<sup>a</sup> and D mutated in pEGFP | This study |
| CS1 | Epn2<sup>160-640</sup> C and D mutated, C<sup>b</sup> in the position of C<sup>a</sup> in pEGFP | This study |
| CF1 | Epn2<sup>160-640</sup> C and D mutated, C<sup>b</sup> and its flanking region in the position of C<sup>a</sup> and its flanking region in pEGFP | This study |
| CS2 | Epn2<sup>160-640</sup> CS and D mutated, C1 in the position of C2 in pEGFP | This study |
| CF2 | Epn2<sup>160-640</sup> Epn2<sup>160-640</sup> C and D mutated, C1 and its flanking region in the position of C2 and its flanking region in pEGFP | This study |
| CDN3-GFP | Epn3 U mutated in pEGFP | This study |
| UDN3-GFP | Epn3 C mutated in pEGFP | This study |
| UCN3-GFP | Epn3 D mutated in pEGFP | This study |
| UCD3-GFP | Epn3 N mutated in pEGFP | This study |
| DN3-GFP | Epn3 U and C mutated in pEGFP | This study |
| CN3-GFP | Epn3 U and D mutated in pEGFP | This study |
| CD3-GFP | Epn3 U and N mutated in pEGFP | This study |
| UN3-GFP | Epn3 C and D mutated in pEGFP | This study |
| UD3-GFP | Epn3 C and N mutated in pEGFP | This study |
| UC3-GFP | Epn3 D and N mutated in pEGFP | This study |
| U3-GFP | Epn3 C, D and N in pEGFP mutated | This study |
| C3-GFP | Epn3 U, D and N mutated in pEGFP | This study |
| D3-GFP | Epn3 U, C and N mutated in pEGFP | This study |
| N3-GFP | Epn3 U, C and D mutated in pEGFP | This study |
| ~~UCDN~~3-GFP | Epn3 U, C, D and N mutated in pEGFP | This study |
| ΔE-CDN3-GFP | Epn3<sup>176-633</sup>U mutated in pEGFP | This study |
| ΔE-UDN3-GFP | Epn3<sup>176-633</sup>C mutated in pEGFP | This study |
| ΔE-UCN3-GFP | Epn3<sup>176-633</sup>D mutated in pEGFP | This study |
| ΔE-UCD3-GFP | Epn3<sup>176-633</sup>N mutated in pEGFP | This study |
| ΔE-DN3-GFP | Epn3<sup>176-633</sup>U and C mutated in pEGFP | This study |
| ΔE-CN3-GFP | Epn3<sup>176-633</sup>U and D mutated in pEGFP | This study |
| ΔE-CD3-GFP | Epn3<sup>176-633</sup>U and N mutated in pEGFP | This study |
| ΔE-UN3-GFP | Epn3<sup>176-633</sup>C and D mutated in pEGFP | This study |
| ΔE-UD3-GFP | Epn3<sup>176-633</sup>C and N mutated in pEGFP | This study |
| ΔE-UC3-GFP | Epn3<sup>176-633</sup>D and N mutated in pEGFP | This study |
| ΔE-U3-GFP | Epn3<sup>176-633</sup>C, D and N mutated in pEGFP | This study |
| ΔE-C3-GFP | Epn3<sup>176-633</sup>U, D and N mutated in pEGFP | This study |
| ΔE-D3-GFP | Epn3<sup>176-633</sup>U, C and N mutated in pEGFP | This study |
| ΔE-N3-GFP | Epn3<sup>176-633</sup>U, C and D mutated in pEGFP | This study |
| ΔE-~~UCDN~~3-GFP | Epn3<sup>176-633</sup>U, C, D and N mutated in pEGFP | This study |

**Suppl Fig. 1.**
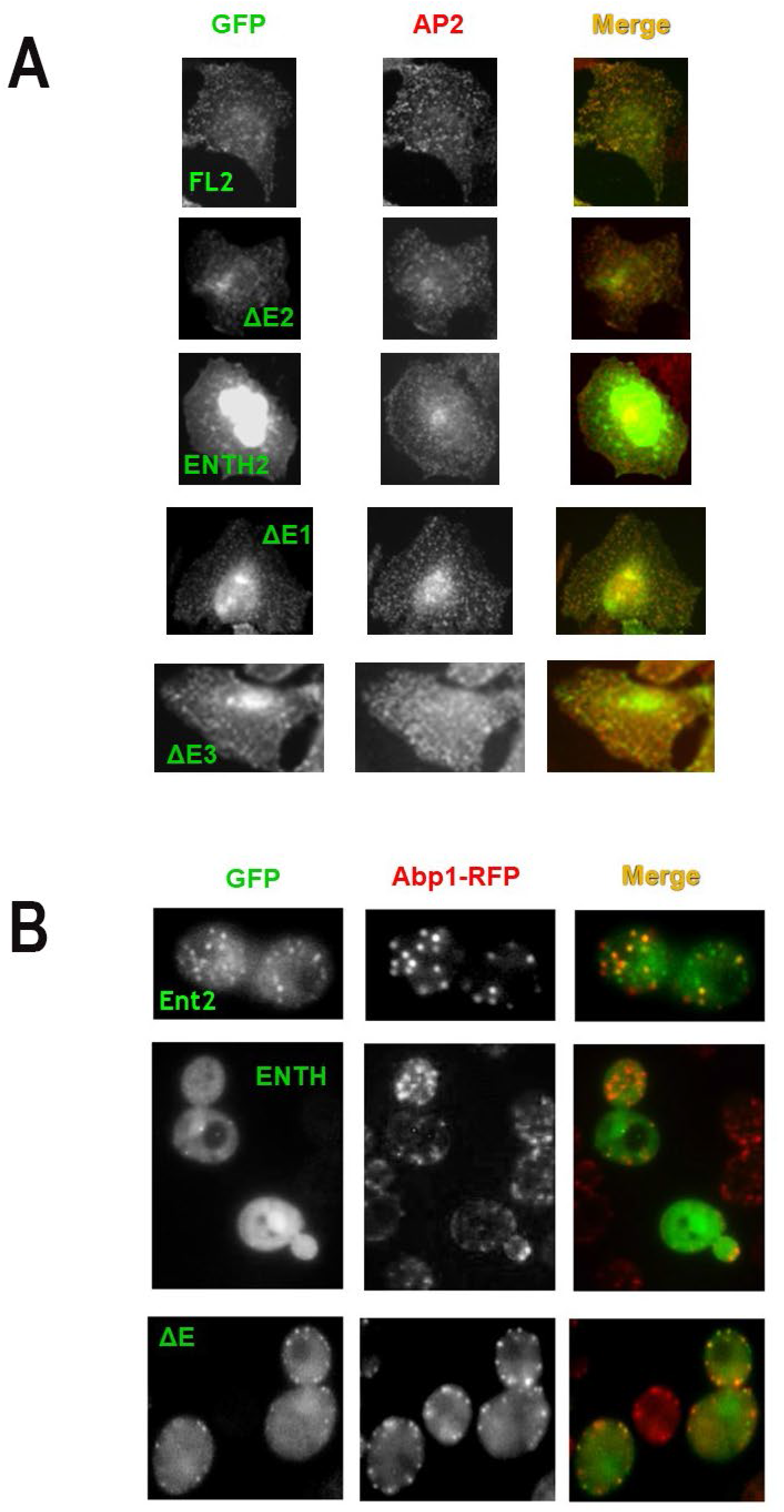

**Suppl Fig. 2.**
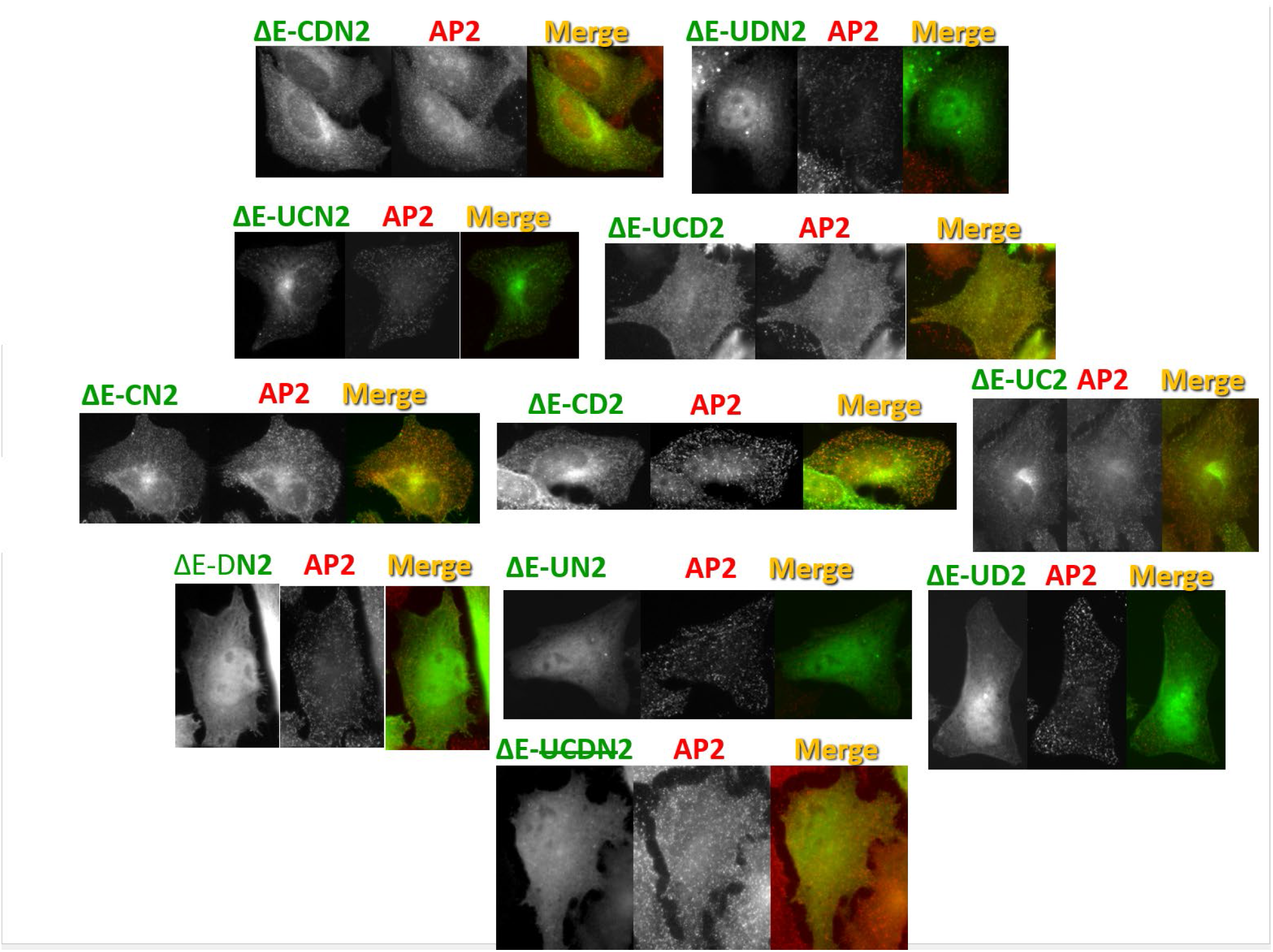

**Suppl Fig. 3.**
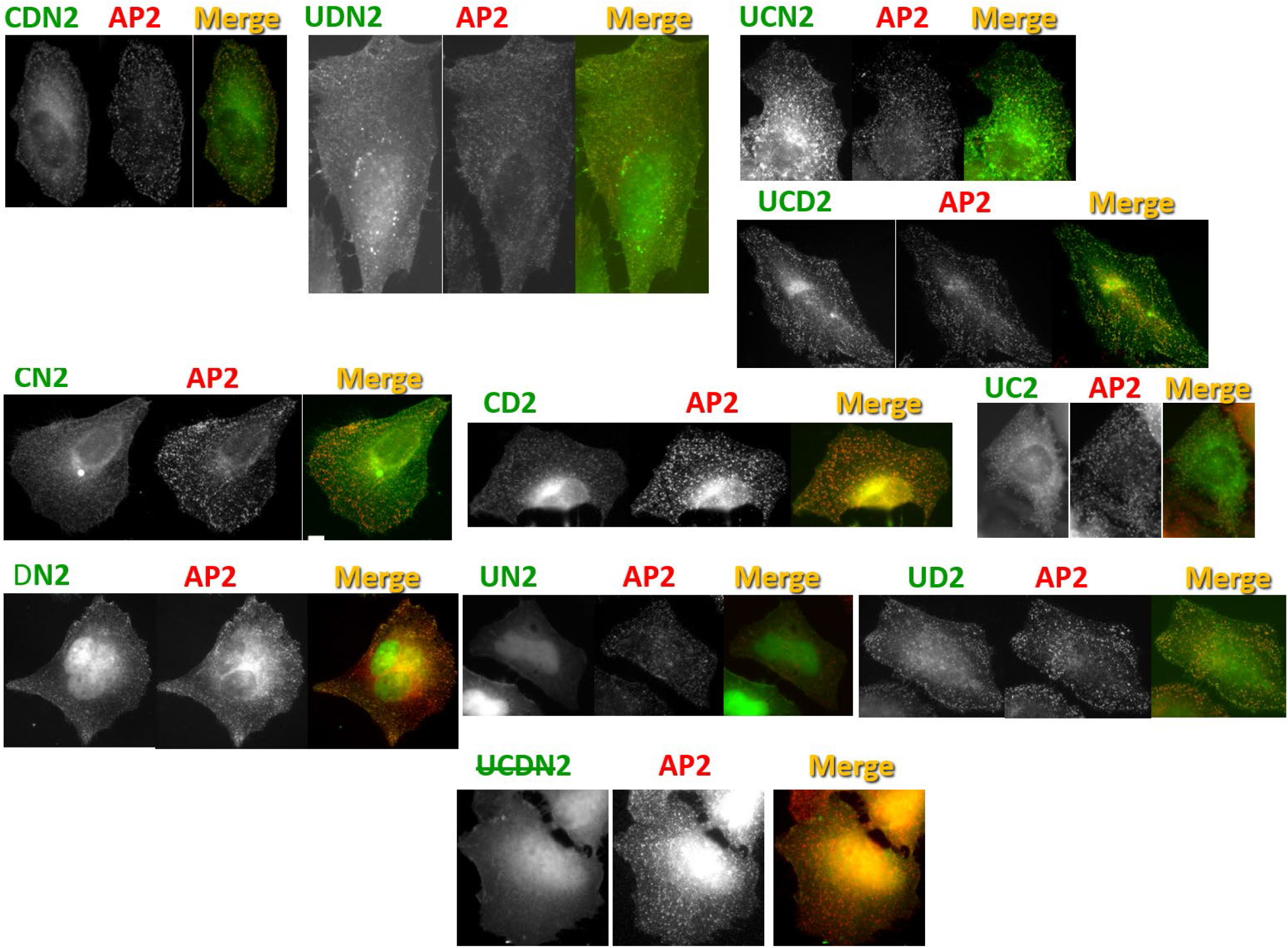

**Suppl Fig. 4.**
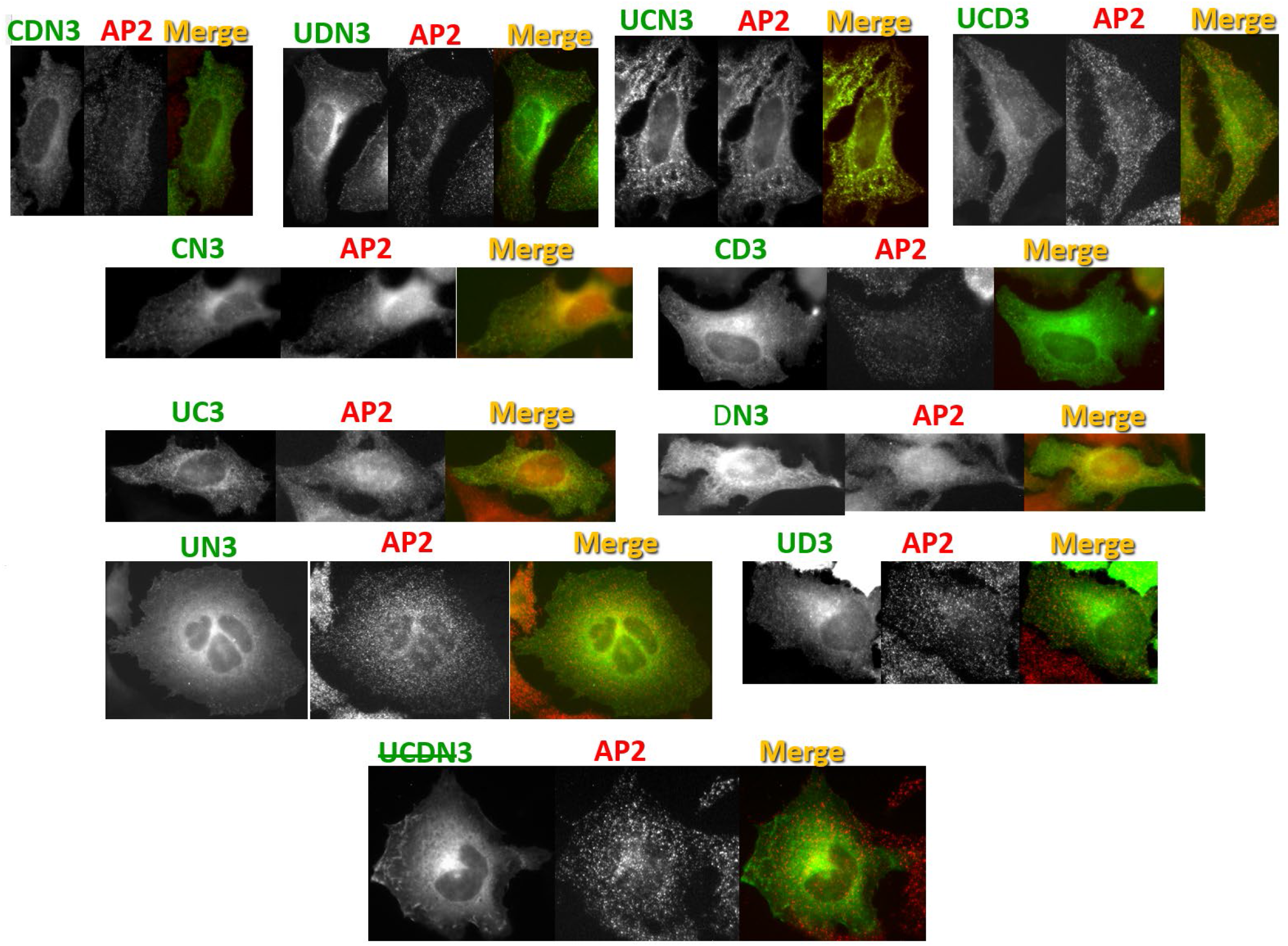

**Suppl Fig. 5.**
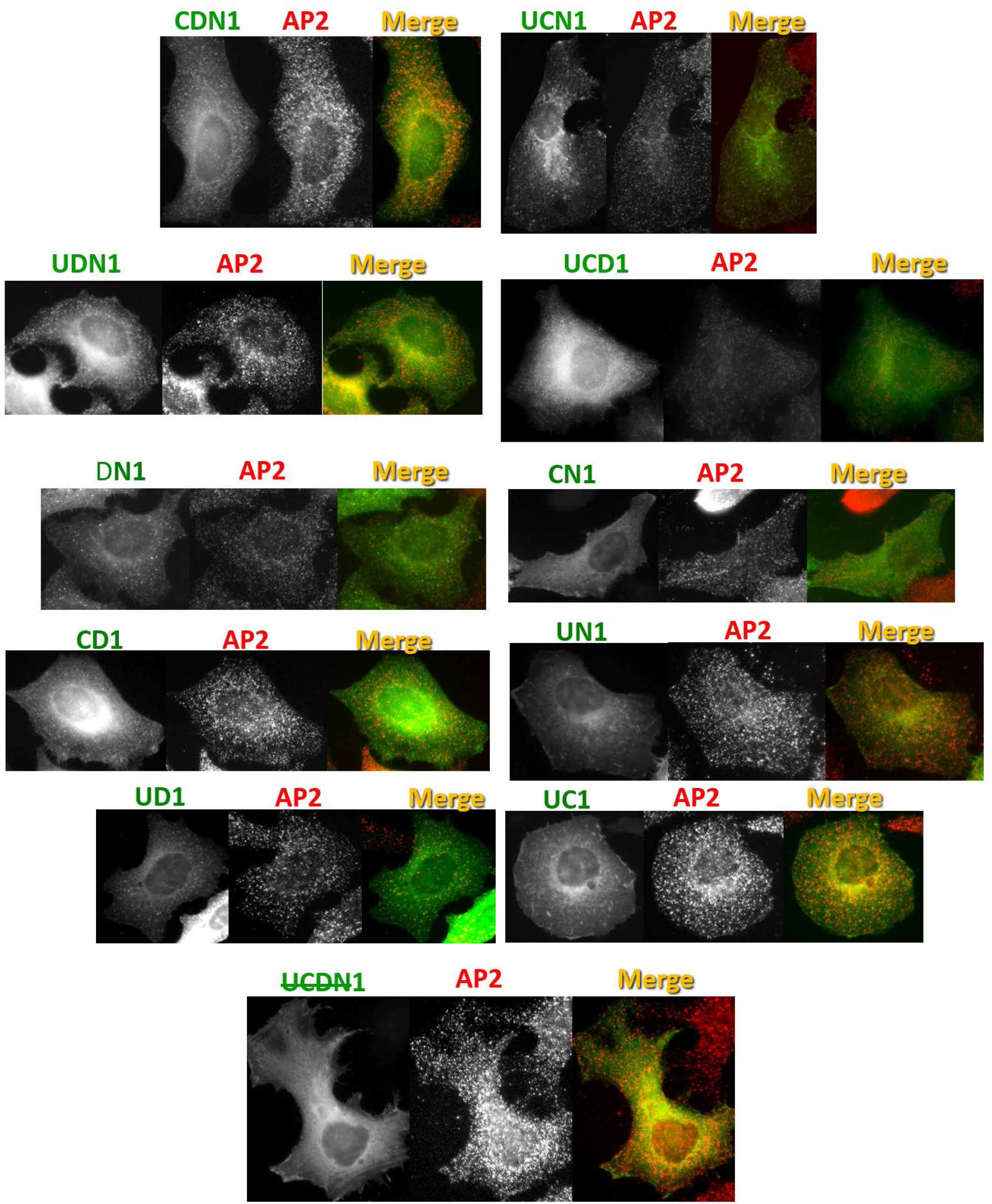

**Suppl Fig. 6.**
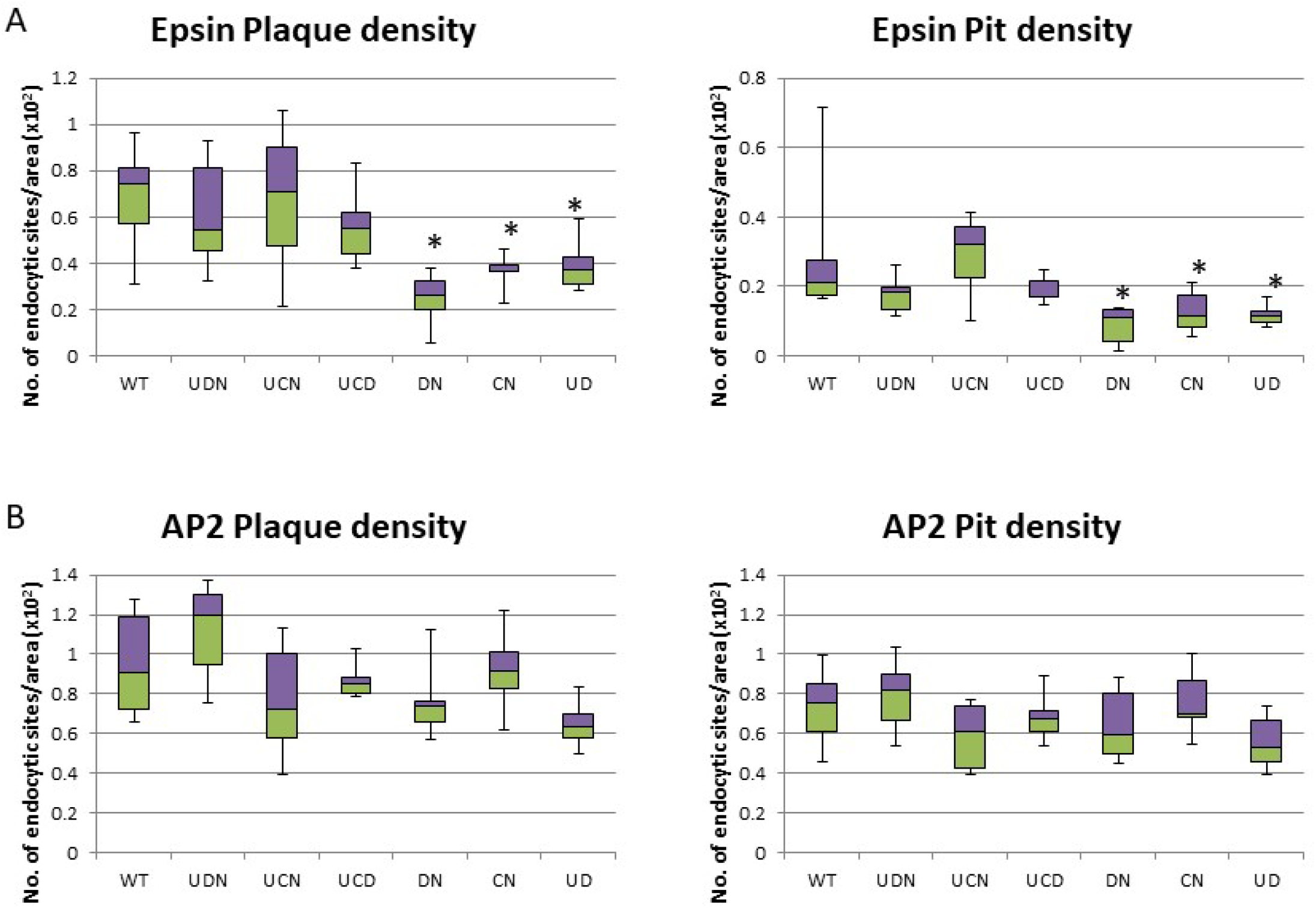

## Notes

### Competing Interest Statement

The authors have declared no competing interest.

